# Disruption of the rice *4-DEOXYOROBANCHOL HYDROXYLASE* unravels specific functions of canonical strigolactones

**DOI:** 10.1101/2023.04.03.535333

**Authors:** Guan-Ting Erica Chen, Jian You Wang, Cristina Votta, Justine Braguy, Muhammad Jamil, Gwendolyn K Kirschner, Valentina Fiorilli, Lamis Berqdar, Aparna Balakrishna, Ikram Blilou, Luisa Lanfranco, Salim Al-Babili

## Abstract

Strigolactones (SLs) regulate many developmental processes, including shoot-branching/tillering, and mediate rhizospheric interactions. SLs are structurally diverse, divided into a canonical and a non-canonical sub-family. To better understand the biological function of particular SLs, we generated CRISPR/Cas9 mutants disrupted in *OsMAX1-1400* or *OsMAX1-1900*, which encode cytochrome P450 enzymes (CYP711A clade) contributing to SL diversity. The disruption of *OsMAX1-1900* did neither affect the SL pattern nor plant architecture, indicating a functional redundancy. In contrast, disruption of OsMAX1-1400 activity, a 4-deoxyorobanchol hydroxylase, led to a complete lack of orobanchol and an accumulation of its precursor 4-deoxyorobanchol (4DO), both of which are a canonical SLs common in different plant species, accompanied by higher levels of the non-canonical methyl 4-oxo-carlactonoate (4-oxo-MeCLA). *Os1400* mutants showed also shorter plant height, panicle and panicle base length, but did not exhibit a tillering phenotype. Hormone quantification and transcriptome analysis revealed elevated auxin levels and changes in the expression of auxin-related, as well as of SL biosynthetic genes. Interestingly, the *Os900/1400* double mutant lacking both orobanchol and 4DO did not show the observed *Os1400* architectural phenotypes, indicating that they are a result of 4DO accumulation. A comparison of the mycorrhization and Striga seed germinating activity of *Os900, Os900/1400*, and *Os1400* loss-of-function mutants demonstrates that the germination activity positively correlates with 4DO content while disrupting *OsMAX1-1400* negatively impact mycorrhizal symbiosis. Taken together, our paper deciphers the biological function of canonical SLs in rice and depicts their particular contributions to establishing architecture and rhizospheric communications.

## Introduction

The apocarotenoid-derived strigolactones (SLs) are a novel class of plant hormones induced under low phosphate (Pi) condition that inhibits shoot branching/tillering (Gomez-Roldan et al., 2008; Umehara et al., 2008) and regulates other plant processes and features, including root development, stem thickness, and leaf senescence (Al-Babili & Bouwmeester, 2015; Fiorilli et al., 2019). Before being recognized as a plant hormone, SLs were first discovered to be the germinating stimulants for root parasitic weeds, such as *Orobanche* and *Striga spp*. (Cook et al., 1966), and later found as an initiation signal for establishing beneficial arbuscular mycorrhizal fungi (AMF) symbiosis, by inducing the AMF hyphal branching (Akiyama et al., 2005; Lanfranco et al., 2018). In addition, SLs can orchestrate plant architecture as SL-deficient plants, such as rice *d17*, consistently exhibit a distinct phenotype comprising higher numbers of branches/tillers, shorter shoot and primary root length when compared to wild-type (WT) (Al-Babili & Bouwmeester, 2015; Gomez-Roldan et al., 2008; Morris et al., 2001), suggesting that SLs are more than rhizospheric signals (Wang et al., 2022a).

SLs are characterized based on their exclusive chemical structure, the lactone ring (D-ring; Fig.1A). The latter is linked to an enol-ether bridge, essential for the SL biological activity (Yoneyama et al., 2018). According to the presence or absence of the BC-ring (Fig.1A), they are classified into canonical and non-canonical SLs, respectively (Al-Babili & Bouwmeester, 2015; Wang et al., 2021). In general, the evolutionarily-conserved SL biosynthetic pathway in land plants starts specifically with 9-*cis*-β- carotene after its isomerization by DWARF27 (D27) (Abuauf et al., 2018). Then, two carotenoid cleavage dioxygenases (CCDs), CCD7 (D17) and CCD8 (D10), cleave successively 9-*cis*-β-carotene into carlactone (CL), the core intermediate of SL biosynthesis *in planta* (Alder et al., 2012; Bruno et al., 2014; Chen et al., 2022; Seto et al., 2014; Wang et al., 2021).

**Figure 1.**
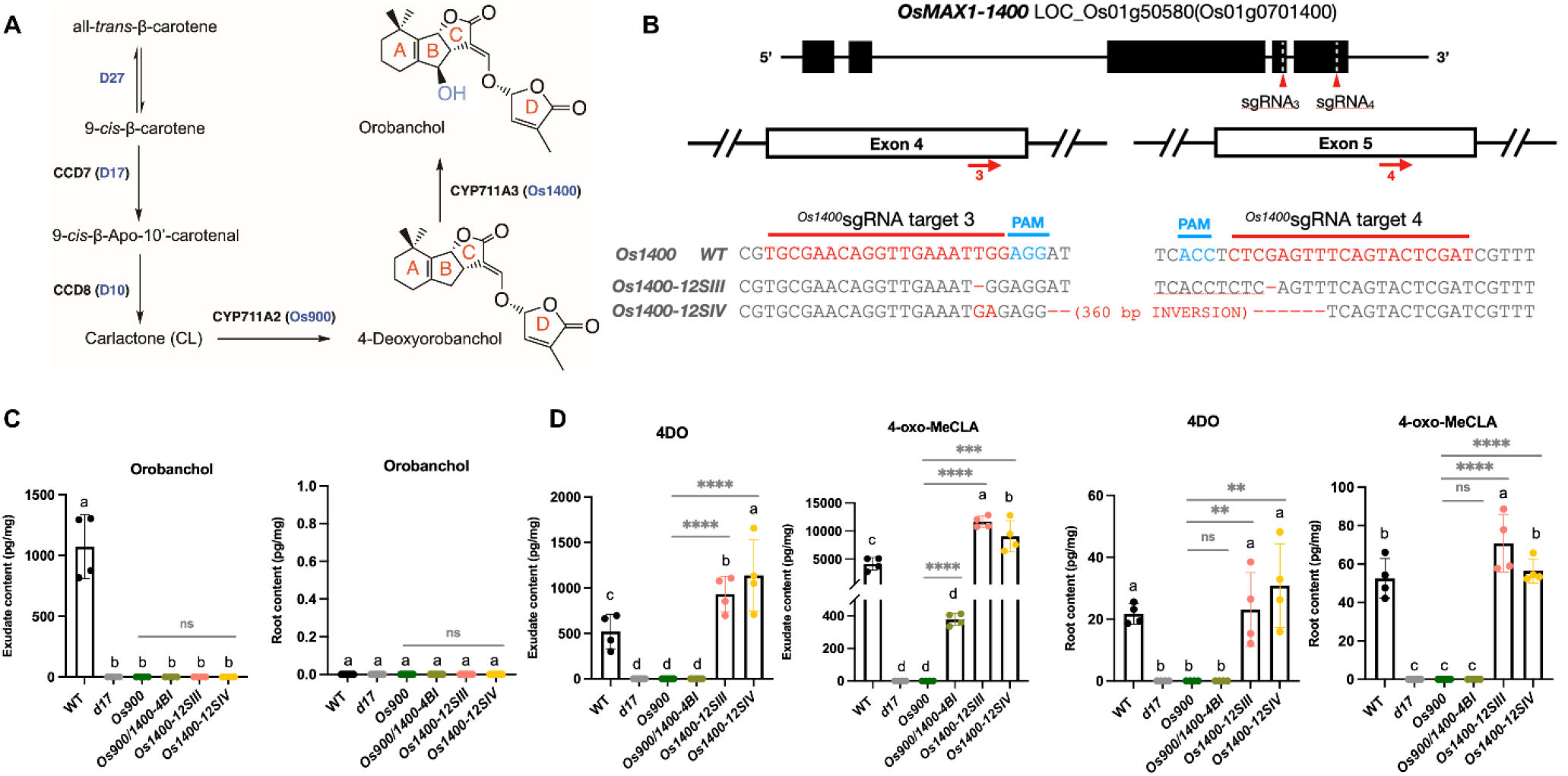
Generation of the *Os1400*-KO lines by CRISPR-Cas9 system. (A) Scheme of the biosynthesis of the rice canonical SLs (the detailed SL biosynthesis pathway depicted in fig. S1. (B) The structure of the *Os1400* gene and the sequences of the two CRISPR-Cas9 target sites indicated by red arrows 3 and 4. Details of the CRISPR-mediated mutations of the two KO lines, *Os1400-12SIII* and *Os1400-12SIV*, are reported in FigS2. (C) Analysis of SLs in root exudates and root tissues of WT, *Os900-KO line, Os900/1400-KO line, Os1400-KO lines, and d17 mutant* grown under constant low-Pi conditions. The data are presented as means ± SD for the number of biological replicates n=4 for (C). Significant values determined by one-way ANOVA are shown with different letter (P < 0.05) when compared to WT, and asterisks indicate statistically significant differences as compared to control by two tailed paired Student’s t-test (*p < 0.05, **p < 0.01; ***p < 0.001; ****p < 0.0001). Abbreviation: 4DO, 4-Deoxyorobanchol

Until now, more than 35 natural SLs, with different chemical structures, are identified *in Plantae* (Yoneyama et al., 2018). Their structural diversity arose from the CL catabolism by MORE AXILLARY GROWTH1 (MAX1) from the cytochrome P450 monooxygenase (CYP) 711A family (Booker et al., 2005; Cardoso et al., 2014; Lazar & Goodman, 2006), and the recently identified CYP722C, CYP712G1, CYP706C37 clades (Li et al., 2023; Wakabayashi et al., 2019; Wang et al., 2022c). In rice, OsMAX1-900 repeatedly oxygenates CL to produce the canonical SL 4-deoxyorobanchol (4DO) *in vivo* (Ito et al., 2022). Based on *in vitro* studies and expression in *Nicotiana benthamiana*, 4DO is further hydroxylated into orobanchol (Oro) by another CYP711A enzyme, OsMAX1-1400, (Zhang et al., 2014).

The biological relevance of such a big family of compounds is yet to be discovered by the scientific community. Therefore, we started our investigation in rice and created CRISPR-mediated mutant lines of *OsMAX1s*. Our previous study surprisingly showed that mutants created by targeting *OSMAX1-900* do not display a typical SL-deficiency phenotype (Ito et al., 2022). This revealed that canonical SLs are not the major tillering regulators, and hence the biological functions of canonical SLs inside plants remain elusive. To have a comprehensive view, in this work we aimed to understand the biological roles of canonical SLs in rice by studying mutants defective of *Osmax1-1400*, the other rice enzyme involved in canonical SLs synthesis.

## Results and Discussion

### Characterization of rice *MAX1* homologs

To investigate the biological functions of canonical SLs in rice, we generated several mutant lines using CRISPR/Cas9 technology. We targeted the biosynthesis of canonical SLs by generating biallelic homozygote *Osmax1-1400* (*Os1400-12SIII* and *-12SIV*, Fig. S2). We also generated *Osmax1-1900* (*Os1900-13DI* and *-13EI*, Fig. S3) rice mutant lines to understand whether the *MAX1-1900* would contribute to SL biosynthesis *in planta*, as MAX1-1900 is phylogenetically from a distinct clade, and only shown to weakly convert CL into CLA *in vitro* (Marzec et al., 2020; Yoneyama et al., 2018).

First, we quantified the known SLs in these mutants’ roots and root exudates, together with *Os900* and SL-deficient *d17*, using Liquid Chromatography Tandem-Mass Spectrometry (LC-MS/MS) under phosphate (Pi) starvation conditions (Fig. 1C; Fig. 1D; Fig. S4). As expected, no SLs were detected in *d17*, while the CRISPR/Cas9-induced deletion impaired the biosynthesis of the canonical SLs, which was evidenced by the absent Oro in the roots and root exudates of *Os1400* mutants (Fig. 1C). This confirmed OsMAX1-1400 as the 4DO hydroxylase *in planta* (Zhang et al., 2014). Interestingly, the accumulation of two SLs, 4DO and the putative methyl 4-oxo-carlactonoate (4-oxo-MeCLA), produced through OsMAX1-900 (Ito et al., 2022), were doubled in the root exudates of *Os1400* when compared to wild-type (WT) plants (Fig. 1D). The production of 4-oxo-MeCLA is likely through an additional step by an uncharacterized methyltransferase (Ito et al., 2022), suggesting that 4DO metabolism is exclusive by OsMAX1-1400. Moreover, the amount of putative non-canonical SLs, CL+30 and oxo-CL (Ito et al., 2022; Wang et al., 2022a), in the root exudates of *Os1400* were comparable to that of WT (Fig. S4), affirming the mutation of *Os1400* mainly affects the canonical SL Oro metabolism.

In contrast, we did not observe remarkable changes in SL composition in the root tissues and root exudates between *Os1900* mutants and WT (Fig. S5), indicating that the role of OsMAX1-1900 might be functionally redundant in SL biosynthesis in plants. This was supported by no consistently significant differences in the shoot and root phenotypes grown under normal or low Pi conditions as well as *Striga* germination activity, when compared to the WT (Fig. S6, Fig. S7).

### Accumulation of 4-deoxyorobanchol negatively regulates rice plant growth and development

Recently, we reported that rice lacking canonical SLs, 4DO and Oro, displays shoot phenotypes comparable to WT, revealing that canonical SLs are not the major players in regulating shoot architectures (Ito et al., 2022). However, rice has duplicated *MAX1* genes during evolution (Marzec et al., 2020); intriguingly, OsMAX1-1400 is the only rice MAX1 acting in the final step of canonical SL biosynthesis (Fig. 1A) (Al-Babili & Bouwmeester, 2015; Zhang et al., 2014). Therefore, we suspected that *Os1400* and its downstream metabolite Oro might hold biological importance in plants. To test this hypothesis, we phenotyped *Os1400, Os900, d17*, and WT rice under soil and hydroponic conditions. Expectedly, the *MAX1* mutants, grown in greenhouse, did not showed high tiller numbers with dwarf appearance, the typical SL-deficit (*d17*) phenotype (Fig. 2A). Instead, the plant height, panicle length, and length of the panicle base (the distance from flag leaf auricle to the panicle base on the primary branch) of *Os1400* mutants were significantly shorter than that of WT and *Os900* (Fig. 2B). No differences were observed in the total number of tillers, productive tillers, and average panicle numbers under greenhouse conditions (Fig. S8). The shorter shoot phenotype was also detected in the hydroponically grown *Os1400* mutants under both normal and low-Pi conditions (Fig. S9; Fig. S10).

**Figure 2.**
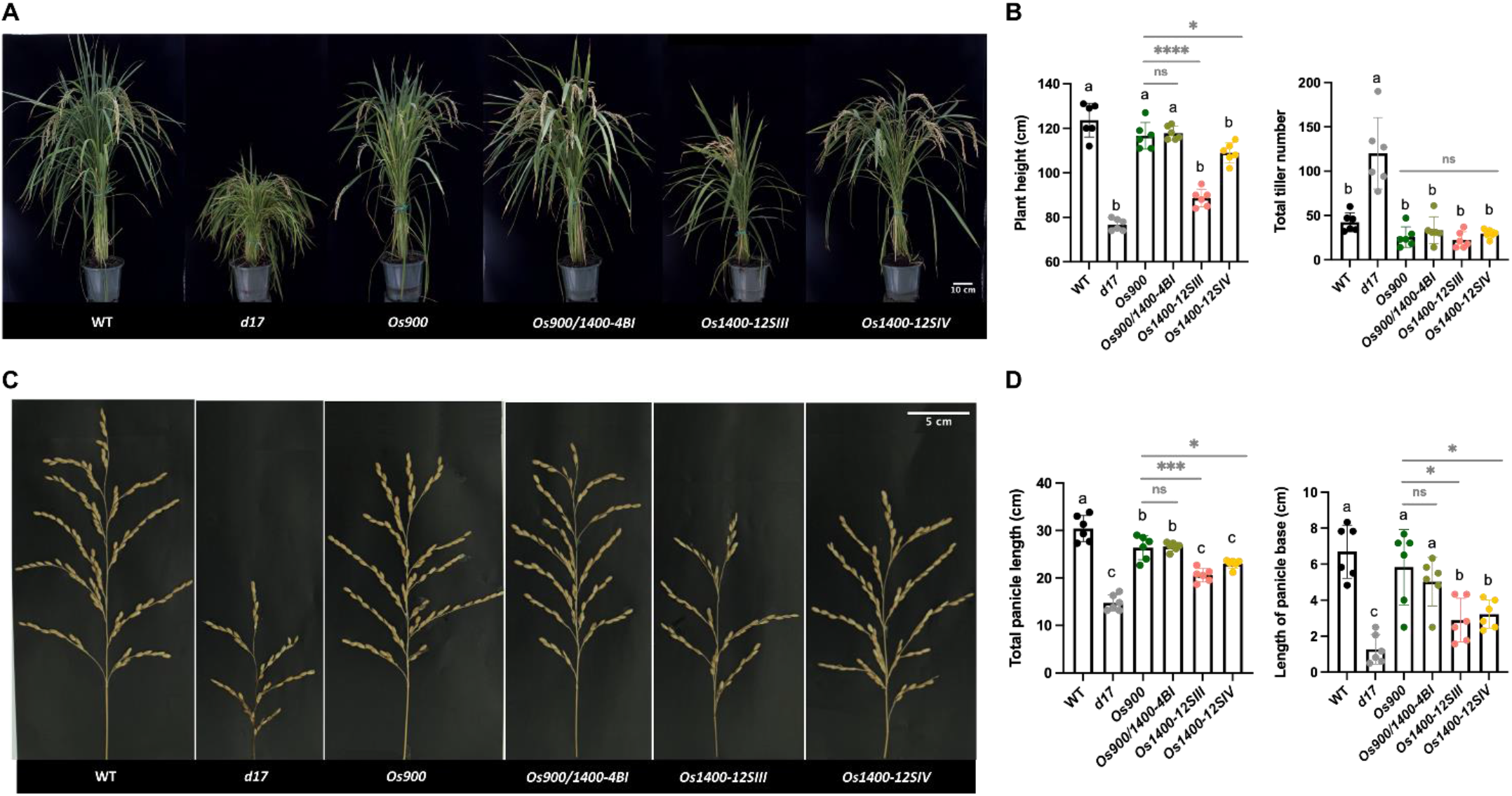
Phenotypic characterization of *OsMAX1* mutants. (A and B) Shoot phenotypes of WT, *Os900*-KO line, Os900/1400-KO line, *Os1400*-KO lines, and d17 mutant grown in soil. Scale bar, 10cm. (C and D) Panicle phenotypes of WT, *Os900*-KO line, *Os900/1400*-KO line, *Os1400*-KO lines, and d17 mutant. Scale bar, 5 cm The data are all presented as means ± SD for the number of biological replicates n=6. Significant values determined by one-way ANOVA are shown with different letter (P < 0.05) when compared to WT, and asterisks indicate statistically significant differences as compared to control by two tailed paired Student’s t-test (*p < 0.05, **p < 0.01; ***p < 0.001; ****p < 0.0001).

It seems that the accumulation of 4DO could be an endogenous unfavorable/toxic signal to rice plants. To confirm the assumption that the shorter shoot phenotype is caused by 4DO accumulation, we exogenously supplied 4DO at 300 nM and 900 nM under normal growth conditions -without triggering SL biosynthesis- to WT and *Os1400* mutants. Interestingly, 300 nM 4DO treatment remarkably increased the difference of shoot length in *Os1400* mutants compared to untreated WT; however, with 900 nM 4DO application, the shoot length in the WT was also suppressed, suggesting that 4DO is a negative regulator of shoot growth and development in rice (Fig. S11). On the other hand, we observed a longer root length tendency in *Os1400* than WT (Fig. S10) under low-Pi conditions, while a decreased tendency in root length and root diameter under normal conditions (Fig. S9; Fig. S12). As SL biosynthesis is triggered by Pi-starvation, we postulated that Pi content might influence the amount of 4DO accumulated in the plant, leading to the different tendency in root length under various growth conditions. Indeed, exogenous 4DO application under normal conditions, mimicking SL biosynthesis, enhanced the root length in all treated rice plants grown in hydroponics (Fig. S11).

Next, to check our rescue hypothesis by reducing the level of 4DO accumulated in *Os1400* mutants, we treated *Os1400* grown under normal conditions with 5 µM TIS108, a MAX1-900 and MAX1-1400 inhibitor (Ito et al., 2022). The root length and crown root numbers of mutants treated with TIS108 were restored to that of the WT (Fig. S13), providing a further evidence that the enzymatic activity on 4DO by OsMAX1-1400 is important for an optimal rice root development. In fact, the root-released 4DO level of *Os1400* mutants was suppressed upon TIS108 treatment, accompanied by the accumulation of the putative non-canonical SLs, CL+30 and oxo-CL (Fig. S14) (Ito et al., 2022; Wang et al., 2022a). Although the shoot length of *Os1400* mutants no longer showed difference to the WT when treated with TIS108, we observed that two-week application of TIS108 unexpectedly decreased the shoot length of WT (Fig. S13), probably due to unspecific compound effects on other CYPs. Hence, to specifically understand the biological roles of canonical SL biosynthesis, we generated a homozygous *Osmax1-900/1400* (*Os900/1400-4BI*) rice double mutant line (Fig. S15), which has full disruption of canonical SLs, to compare with the TIS108 observation. Notably, the observed phenotypes of *Os1400* mutants were no longer present in the double mutant in all growth conditions (Fig. 2; Fig. S8; Fig. S9; Fig. S10: Fig. S12), together with the absence of canonical SLs and the accumulation of non-canonical SLs in the roots and root exudates of *Os900/1400* (Fig. 1C; Fig. 1D; Fig. S4). Sharing the same rescue mechanism as TIS treatment, suppressing both 4DO and Oro biosynthesis, *Os900/1400* provides a stronger genetic evidence without the possibilities of chemical effects. These suggest that the metabolism of 4DO by OsMAX1-1400 is required for normal rice growth and development.

One previous study had contradictory results, showing a high-tillering phenotype with low canonical SLs in the Bala rice that has both a deletion of *MAX1-900* and *MAX1-1400* (Cardoso et al., 2014). However, our observations hold stronger genetic evidence since these genetic materials were compared within the same cultivar, while the tillering phenotype of Bala rice might be a consequence of other genetic regulations. Notably, 4-oxo-MeCLA was present in the root exudate of *Os900/1400*, as well as in Bala rice root exudate (Cardoso et al., 2014), indicating that the biosynthesis of this non-canonical SL is independent of MAX1-900 and MAX1-1400. Overall, we can conclude that 4DO and Oro are not tillering-inhibitory regulators, but their unbalanced metabolism negatively affects the physiological development of rice plants.

Furthermore, we investigated the rice transcriptome of the *Os1400* grown under normal and low-Pi conditions compared to WT by RNAseq (Data S1). In the differentially expressed genes (DEGs), there were 1712 upregulated genes and 1465 downregulated genes under normal conditions, while 5890 upregulated genes and 3569 downregulated genes were observed under low-Pi conditions (Fig. S16). None of the DEGs related to tillering or SL biosynthesis was observed under normal conditions (Tables S1); in contrast, the transcripts of SL biosynthesis and signaling were generally downregulated under low-Pi conditions (Fig. 3A, Tables S2), suggesting that the 4DO accumulation might lead to a negative feedback loop in *Os1400* mutants. Interestingly, under both conditions, we observed many downregulated auxin-related genes in *Os1400* mutants (Fig. 3B, Tables S1-S2).

**Figure 3.**
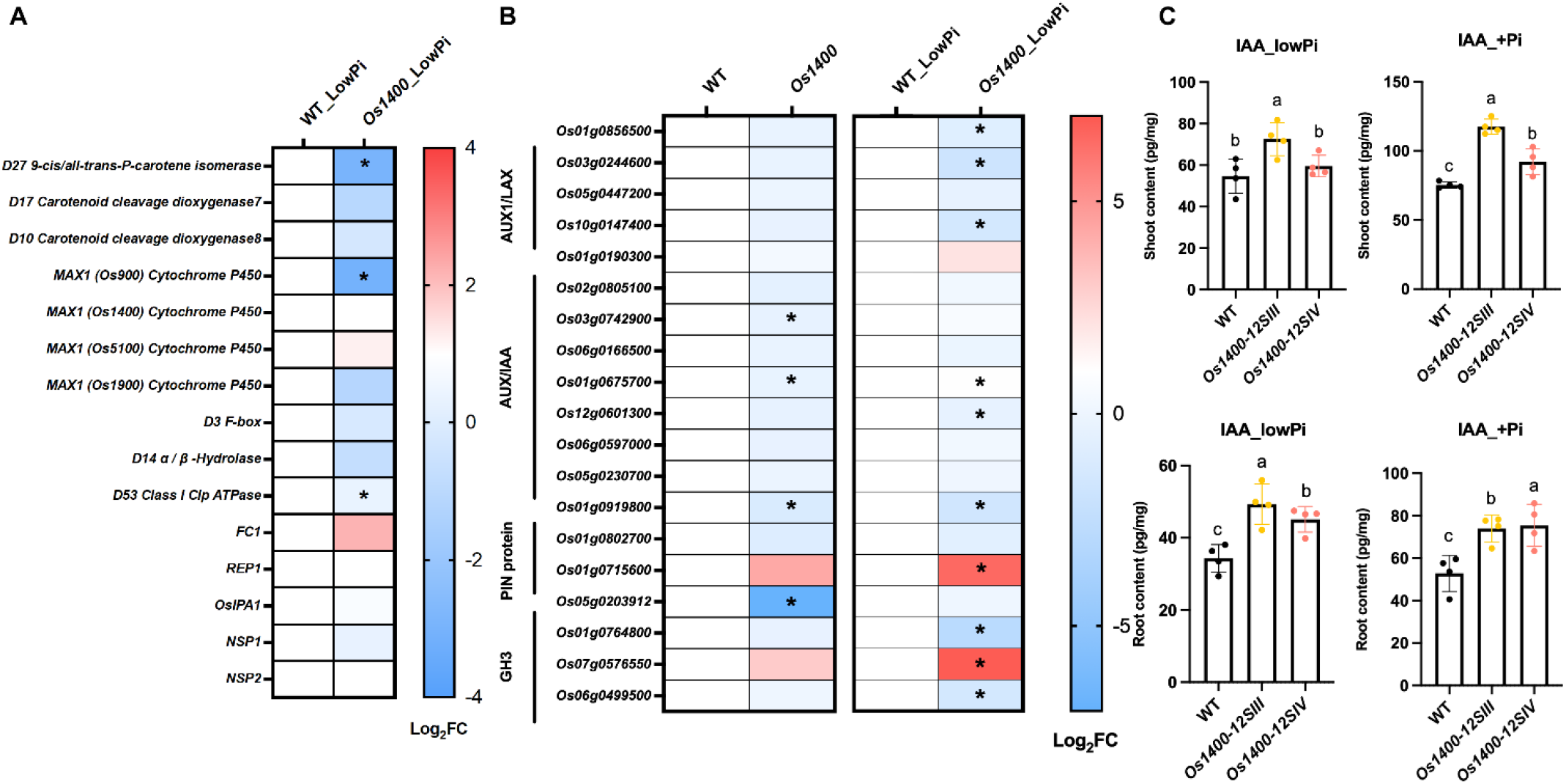
Transcriptome and hormone analysis of *Os1400*-KO lines. (A) Differentially expressed genes (DEGs) of SL biosynthesis and signaling related gene. Expression pattern was shown in log2FoldChange (Log2FC). Statistically significant differences indicated by adjusted p-value (*< 0.05). (B) Heat map analysis of DEGs involved Auxin pathways. AUXIN1/LIKE-AUX1 (AUX1/LAX) are major auxin influx carriers; AUXIN/indole-3-acetic acid (AUX/IAA) are transcriptional repressors; the PIN-FORMED (PIN) proteins are secondary transporters in the efflux of auxin; GH3 gene family encodes auxin-amido synthetases. Expression pattern was shown in log2FoldChange (Log2FC). Statistically significant differences indicated by adjusted p-value (*< 0.05). (C) Analysis of IAA (auxin) in root and shoot of WT, *and Os1400*-KO *lines* grown under constant low-Pi and +Pi conditions. Abbreviation: IAA, indole-3-acetic acid. The data are all presented as means ± SD for the number of biological replicates n=3 for (A and B), and n=4 for (C). Significant values determined by one-way ANOVA are shown with different letter (P < 0.05) when compared to WT, and asterisks indicate statistically significant differences as compared to control by two tailed paired Student’s t-test (*p < 0.05, **p < 0.01; ***p < 0.001; ****p < 0.0001).

Auxin regulates meristem activities and interplays with SLs on the root and shoot development (Su et al., 2011; Xiao et al., 2019; Yang et al., 2019); thus, we hypothesized the phenotypes observed in *Os1400* mutants could be linked to auxin. We then determined the hormone content of auxin (IAA), gibberellin (GA), abscisic acid (ABA), salicylic acid (SA), and jasmonic acid (JA) in roots and shoot bases (root-shoot junction) of *Os1400* mutants under normal and low Pi conditions. We did not detect any consistent significant difference in the levels of GAs, ABA, SA, or JA (Fig. S17), but observed a remarkable increase of IAA level in root and shoot bases in both growth conditions, compared to the WT (Fig. 3C; Fig. S17). Consistently, excess of IAA content, by either overexpression of *OsPIN2* or exogenous IAA application, causes shorter shoot phenotype in rice (Liu et al., 2019; Sun et al., 2019). Furthermore, we performed 5-Ethynyl-2’-deoxyuridine (EdU) staining to visualize proliferating cells and measured the root meristem length. Yet, no clear differences were observed between *Os1400* mutants and WT (Fig. S18). Accordingly, we demonstrated that the shoot phenotypes of *Os1400* plants are likely due to auxin homeostasis modulation, but the role of auxin in *Os1400* roots needs further investigation.

### The metabolism of rice canonical SL, 4-deoxyorobanchol, is associated to rhizospheric signals

Although all SLs seem to be communicating signals in the rhizosphere (Ito et al., 2022; Wang et al., 2022a), their bioactivities - to induce AM hyphal branching and trigger parasitic seed germination - depend largely on their chemical structures (Gobena et al., 2017; Mori et al., 2016). The distinguishable root-released SL compositions in *Os900, Os900*/*1400*, and *Os1400* (Fig. 1; Fig. S1) make them good candidates to investigate the possible functions of these metabolites in the rhizosphere. We then tested the germination activity of *Os900, Os1400*, and *Os900*/*1400* root exudates on *Striga hermonthica* seeds. Compared to WT exudates, we observed more than 40% decrease in the *Striga* germination of *Os900* exudates and an even lower germination rate of *Os900/1400* exudates (Fig. 4A), indicating that 4-oxo-MeCLA is not a predominate germinating signal for *Striga* (Fig. S4). Although the stimulation activity on *Striga* germination of *Os1400* exudates was comparable to the WT at 1:1 dilution, we found an increased tendency at 1:3 dilution (Fig. 4A; Fig. S19A). This reveals that 4DO is a stronger germination cue for *Striga* seeds than Oro, supported experimentally by 10 µM Oro exerting less seed germination activity than 1 µM *rac*-GR24, a 4DO-like SL analog (Fig. S19B).

**Figure 4.**
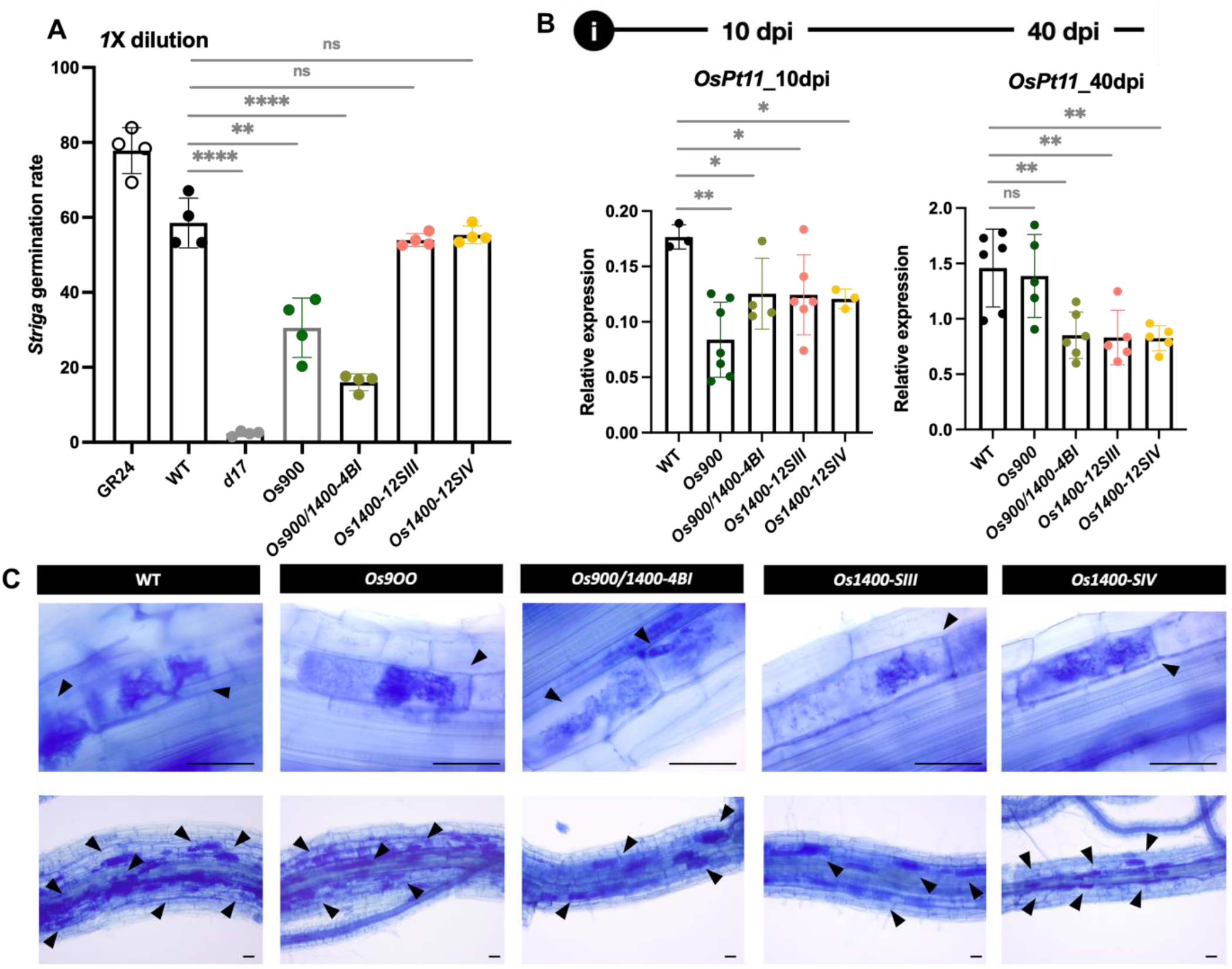
Assessment of rhizospheric interactions. Effect of *Os1400*-KO lines on (A) the germination of root parasitic weed *Striga* and (B and C) the arbuscule formation.. The *R. irregularis* colonization was quantified by measuring the expression of an AM marker gene (*OsPT11*) (B). Arbuscule formation at 10 dpi and 40 dpi. Arrows indicate arbuscule containing cells. Scale bars, 50 µm (C). The data are all presented as means ± SD for the number of biological replicates n=4 for (A), and n≥3 for (B). Significant values determined by one-way ANOVA are shown with different letter (P < 0.05) when compared to WT, and asterisks indicate statistically significant differences as compared to control by two tailed paired Student’s t-test (*p < 0.05, **p < 0.01; ***p < 0.001; ****p < 0.0001).

Moreover, Oro seems to be the preferable signal for AM symbiosis (Mori et al., 2016), we then investigated the role of 4DO and Oro in establishing AM symbiosis by comparing *Os900, Os1400, Os900*/1400, and WT. For this purpose, we examined the root colonization with the AMF *Rhizophagus irregularis* after 10- and 40-days post inoculation (dpi) and measured the transcript level of *OsPT11*, a specific AM inducible Pi-transporter gene (Güimil et al., 2005). At 10 dpi, there was a delay in colonization of all mutant roots compared to WT roots; whereas, at 40 dpi, the colonization of *Os1400* and *Os900*/1400 mutants was surprisingly much lower than that of the WT and *Os900* (Fig. 4B; Fig. S20), which indicates that *MAX1-1400* is crucial for maintaining AM colonization. Additionally, absence of *MAX1-1400* did not influence the intraradical fungal structures of the arbuscules. Instead, they appeared well developed and regularly branched (Fig. 4C), suggesting that *MAX1-1400* does not affect the fungal morphology. Therefore, we can speculate that *MAX1* duplication is highly associated with AM symbiosis; besides, the decreased colonization level of *Os900*/1400 mutant further supported that the reduced AM symbiotic pattern upon TIS108 application in WT plants (Ito et al., 2022) might be a consequence of *MAX1-1400* suppression.

Taken together, the generation of the CRISPR-mediated *Os1400* rice mutants allowed us to reveal the biological roles of canonical SLs in plant growth and development, and as well confirmed OsMAX1-1400 as the 4DO hydroxylase *in planta*. Finally, we can conclude that the canonical SLs, 4DO and Oro, are critical rhizospheric signals for the interaction with AMF and root parasitic plants, and the *MAX1* duplication might be also evolutionary required for beneficial symbiosis in rice. Importantly, knocking out entire canonical SLs, without damaging plant architecture in the *Os900/1400* double mutant, seems promising to reduce the yield loss caused by *Striga* and other root parasitic plants; thus, offering an alternative way to improve global food security.

## Material and Methods

### Plant material and growth conditions

*Oryza sativa* Nipponbare *d17* (Butt et al., 2018), *max1-900* (Ito et al., 2022), *max1-1400, max1-900/1400, max1-1900*, and WT rice plants were grown under controlled conditions (a 12 h photoperiod, 200-µmol photons m^−2^ s^−1^ and day/night temperature of 27/25 °C). All rice seeds were first surface-sterilized in a 50% sodium hypochlorite solution with 0.01 % Tween-20 for 15 min, then rinsed with sterile water, before being germinated in the dark overnight. The pre-germinated seeds were placed on Petri dishes containing half-strength liquid Murashige and Skoog (MS) medium and incubated in a growth chamber for 7 days. Thereafter, the seedlings were transferred into 50mL black falcon tubes filled with half-strength modified Hoagland nutrient solution with adjusted pH to 5.8. The nutrient solution consisted of 5.6 mM NH_4_NO_3_, 0.8 mM MgSO_4_·7H_2_O, 0.8 mM K_2_SO_4_, 0.18 mM FeSO_4_·7H_2_O, 0.18 mM Na_2_EDTA·2H_2_O, 1.6 mM CaCl_2_·2H_2_O, 0.8 mM KNO_3_, 0.023 mM H_3_BO_3_, 0.0045 mM MnCl_2_·4H_2_O, 0.0003 mM CuSO_4_·5H_2_O, 0.0015 mM ZnCl_2_, 0.0001 mM Na_2_MoO_4_·2H_2_O and 0.4 mM K_2_HPO_4_·2H_2_O.

### Generation of *Os1900, Os1400*, and *Os1400/900* plants

Two guide RNAs [gRNAs; single gRNA3 (sgRNA3), 5’ -tgcgaacaggttgaaattgg-3’ and sgRNA4, 5’ -ctcgagtttcagtactcgat-3’] were designed to target the rice (O. sativa L. ssp. japonica cv. Nipponbare) *OsMAX1-1400* (Os01g0701400 /AP014957) gene. By using Golden Gate cloning, the tRNA-gRNA-Cas9 cassette was assembly into the pRGEB32 binary vector that has hygromycin resistance gene for selection. With mature seeds, Nipponbare calli were induced and transformed with *Agrobacterium tumefaciens* EHA105 culture containing the plasmid of interest. Later, shoots and roots were regenerated in a Percival growth chamber (CLF Plant Climatics GmbH, model CU 36L5), and then transferred to soil and grown in a greenhouse at 28°C day/22°C night.

Genomic DNA was extracted from the rice young leaves, and plant transgenicity and mutagenicity were demonstrated. Through polymerase chain reaction (PCR) amplification, the transgenic plants were recognized when the pRGEB32-specific primers, pRGEB32-F (5′-ccacgtgatgtgaagaagtaagataaactg-3′) and pRGEB32-R (5′-gataggtttaagggtgatccaaattgagac-3′), bind to the surrounding region of the insertion sites in the pRGEB32 vector. For identifying CRISPR-mediated mutations, the DNA region that includes the sgRNA target sites were amplified using genome specific primers *Os1400* sg3-sg4 F (5’-tcagcgcgctcacttacga -3’) and *Os1400* sg4 F1 (5’-atcccaagaacttcccggag-3’).

### Hydroponic culture of rice seedlings

The hydroponic culture system is built with 50-mL black falcon tubes with punctured caps inserted with a 1.5-ml bottomless Eppendorf tube in the center. Nutrition solution, containing normal (+Pi) or low 0.004 mM K_2_HPO_3_·3H_2_O (lowPi), was applied to the transferred 1-week old seedlings for the following 2 weeks. The solutions were changed every 3 days, and adjusted to pH 5.8 every time before applying. All plants were kept in the solution for 3 weeks, except the plants for 4DO application and EdU staining were 10-days seedlings.

### Phenotyping in pots under greenhouse conditions

To study the phenotype of *Os1400* and *Os900/1400* mutants, seedlings were transferred into pots packed with soil. The soil were soaked with half-strength modified Hoagland nutrient solution in advance. The nutrient solution comprised 5.6 mM NH_4_NO_3_, 0.8 mM MgSO4.7H_2_O, 0.8 mM K_2_SO4, 0.18 mM FeSO_4_.7H_2_O, 0.18 mM Na_2_EDTA.2H_2_O, 1.6 mM CaCl_2_.2H_2_O, 0.8 mM KNO_3_, 0.023 mM H_3_BO_3_, 0.0045 mM MnCl2.4H_2_O, 0.0003 mM CuSO_4_.5H2O, 0.0015 mM ZnCl_2_, 0.0001 mM Na2MoO4.2H_2_O, and 0.4 mM K_2_HPO_4_.2H_2_O. The pH of the solution was adjusted to 5.8, and the solution was applied every third day. On day 120, phenotypic data were recorded. The plants were grown in a greenhouse from February to May 2022, in Thuwal (Saudi Arabia).

### Exogenous applications of 4DO and TIS108

For investigating the effect of 4DO (Olchemim, Czech Republic) on different genotypes, 1-week-old seedlings were grown hydroponically in half-strength Hoagland nutrient solution containing 0.4 mM K_2_HPO_4_·2H_2_O (+Pi), 300 nM or 900 nM 4DO (dissolved in acetone), or the corresponding volume of the solvent (mock; acetone) for 10 or 14 days. The solution was changed three times per week, adding the chemical at each renewal.

For investigating the effect of TIS108, 2-week-old rice seedlings were grown hydroponically in half-strength Hoagland nutrient solution containing 0.4 mM K_2_HPO_4_·2H_2_O (+Pi), 5 µM TIS108 (dissolved in acetone), or the corresponding volume of the solvent (mock; acetone) for 14 days. The solution was changed twice per week, adding the chemical at each renewal.

### SL quantification in root tissues and exudates

Analysis of SLs in rice root exudates was performed according to the published protocol (Wang et al., 2022b). Briefly, root exudates spiked with 2 ng of GR24 were brought on a C_18_-Fast Reversed-SPE column (500 mg/3 mL), preconditioned with 3 mL of methanol and followed with 3 mL of water. After washing with 3 mL of water, SLs were eluted with 5 mL of acetone. Thereafter, SLs-containing fraction was concentrated to SL aqueous solution (∼500 μL), followed by 1 mL of ethyl acetate extraction. 750 μL of SL enriched fraction was dried under vacuum. The final extract was re-dissolved in 100 μL of acetonitrile: water (25:75, v:v) and filtered through a 0.22 μm filter for LC-MS/MS analysis.

SL extraction from root tissues was followed the procedure (Wang et al., 2019). Around 25 mg of lyophilized and grinded rice root tissues, spiked with 2 ng of GR24, were extracted twice with 2 mL of ethyl acetate in an ultrasound bath (Branson 3510 ultrasonic bath) for 15 min, followed by centrifugation for 8 min at 3800 rpm at 4 °C. The two supernatants were combined and dried under vacuum. The residue was dissolved in 50 μL of ethyl acetate and 2 mL of normal hexane, purifying with a Silica Cartridges SPE column (500 mg/3 mL). After washing with 3 mL of hexane, SLs were eluted in 3 mL of ethyl acetate and evaporated to dryness under vacuum. The final extract was re-dissolved in 150 μL of acetonitrile: water (25:75, v:v) and filtered through a 0.22 μm filter for LC-MS/MS analysis.

SLs were quantified by LC-MS/MS using UHPLC-Triple-Stage Quadrupole Mass Spectrometer (Thermo Scientific™ Altis™). Chromatographic separation was achieved on the Hypersil GOLD C_18_ Selectivity HPLC Columns (150 × 4.6 mm; 3 μm; Thermo Scientific™) with mobile phases consisting of water (A) and acetonitrile (B), both containing 0.1% formic acid, and the following linear gradient (flow rate, 0.5 mL/min): 0– 15 min, 25%–100 % B, followed by washing with 100 % B and equilibration with 25 % B for 3 min. The injection volume was 10 μL, and the column temperature was maintained at 35 °C for each run. The MS parameters of Thermo ScientificTM Altis™ were as follows: positive ion mode, ion source of H-ESI, ion spray voltage of 5000 V, sheath gas of 40 arbitrary units, aux gas of 15 arbitrary units, sweep gas of 20 arbitrary units, ion transfer tube gas temperature of 350 °C, vaporizer temperature of 350 °C, collision energy of 17 eV, CID gas of 2 mTorr, and full width at half maximum (FWHM) 0.2 Da of Q1/Q3 mass. The characteristic Multiple Reaction Monitoring (MRM) transitions (precursor ion → product ion) were 331.15→216.0, 331.15→234.1, 331.15→97.02 for 4-deoxyorobanchol; 347.14→329.14, 347.14→233.12, 347.14→ 205.12, 347.14→97.02 for orobanchol; 361.16→ 247.12, 361.16→177.05, 361.16→208.07, 361.16→97.02 for putative 4-oxo-MeCLA; 333.17→219.2, 333.17→173.2, 333.17→201.2, 333.17→97.02 for putative 4-oxo-hydroxyl-CL (CL+30); 317.17→164.08, 317.17→97.02 for putative oxo-CL (CL+14); 299.09→158.06, 299.09→157.06, 299.09→97.02 for GR24.

### Quantification of plant hormones

Quantification of endogenous hormones was followed the procedure (Wang et al., 2021). Briefly, 15 mg freeze-dried ground root or shoot base tissues were spiked with internal standards D6-ABA (10 ng), D2-GA1 (10 ng), D2-IAA (10 ng), D4-SA (10 ng), and D2-JA (10 ng) along with 750 µL of methanol. The mixture was sonicated for 15 min in an ultrasonic bath (Branson 3510 ultrasonic bath), followed by centrifugation for 5 min at 14,000 × g at 4 °C. The supernatant was collected, and the pellet was re-extracted with 750 µL of the same solvent. Then, the two supernatants were combined and dried under a vacuum. The sample was re-dissolved in 100 μL of acetonitrile:water (25:75, v-v) and filtered through a 0.22 μm filter for LC–MS analysis.

Plant hormones were analyzed using LC-MS/MS using UHPLC-Triple-Stage Quadrupole Mass Spectrometer (Thermo Scientific™ Altis™). Chromatographic separation was achieved on the Hypersil GOLD C_18_ Selectivity HPLC Columns (150 × 4.6 mm; 3 μm; Thermo Scientific™) with mobile phases consisting of water (A) and acetonitrile (B), both containing 0.1% formic acid, and the following linear gradient (flow rate, 0.5 mL/min): 0–10 min, 15%–100 % B, followed by washing with 100 % B for 5 min and equilibration with 15 % B for 2 min. The injection volume was 10 μL, and the column temperature was maintained at 35 °C for each run. The MS parameters of Thermo ScientificTM Altis™ were as follows: positive ion mode for IAA and negative mode for GA, ABA, SA, and JA, ion source of H-ESI, ion spray voltage of 3000 V, sheath gas of 40 arbitrary units, aux gas of 15 arbitrary units, sweep gas of 0 arbitrary units, ion transfer tube gas temperature of 350 °C, vaporizer temperature of 350 °C, collision energy of 20 eV, CID gas of 2 mTorr, and full width at half maximum (FWHM) 0.4 Da of Q1/Q3 mass. The characteristic Multiple Reaction Monitoring (MRM) transitions (precursor ion → product ion) were The characteristic MRM transitions (precursor ion → product ion) were 176.2 → 130 for IAA; 263.2 → 153.1, 263.3 → 204.1, 263.3 → 219.1 for ABA; 347.2 → 259.1, 347.2 → 273 for GA1; 345.1 → 143, 345.1 → 239 for GA3; 137.1 → 93.15, 137.1 → 65.1 for SA; 209.15 → 59.05, 209.15 → 93.04 for JA; 178.2 → 132 for D2-IAA; 269.2 → 159.1 for D6-ABA; 349.1 → 261.1 for D2-GA1; 141.0 → 97.0 for D4-SA ; 211.0 → 61.0 for D2-JA.

### *Striga hermonthica* seed germination bioassays

*Striga* seed germination bioassay was carried out based on the protocol (Jamil et al., 2012; Wang et al., 2022). Briefly, 10-day-old pre-conditioning *Striga* seeds were supplied with 50 μL of extracted root exudates of different rice genotypes. After application, *Striga* seeds were incubated at 30 °C in the dark for 24 hours. Germinated (seeds with radicle) and non-germinated seeds were counted under a binocular microscope to calculate germination rate (%) by using SeedQuant software (Braguy et al., 2021).

### RNA library preparation and transcriptomic analysis

Total rice root RNA was extracted with TRIzol™ (Invitrogen, https://www.thermofisher.com/de/de/home.htmL) using a Direct-zol RNA Miniprep Plus Kit following the manufacturer’s instructions (ZYMO RESEARCH; USA). RNA quality was checked with a Agilent 2100 Bioanalyzer, and RNA concentration was measured using a Qubit 3.0 Fluorometer. The cDNA libraries were constructed following standard protocols and paired-end sequenced on an Illumina NextSeq Sequencer (Illumina HiSeq 4000) by Novogene Bioinformatics Technology Co., Ltd. Total reads were mapped to the rice transcripts using HISAT2 (Kim et al., 2019). Differential gene expression was examined using DESeq2 and established by false discovery rate (FDR) ≤ 0.05 (Love et al., 2014).

### Plant material and growth conditions for *R. irregularis* root colonization

Seed of wild type (cultivar Nipponbare) and independent lines of four *Osmax1* - rice mutants (*Os900, Os900/1400, Os1400-12SIII and -12SIV)* were germinated in pots containing sand and incubated for ten days in a growth chamber under a 14-h light (23 °C)/10-h dark (21 °C). Plants were inoculated with ∼1000 sterile spores of *Rhizophagus irregularis* DAOM 197198 (Agronutrition, Labège, France). Plants were grown in sterile quartz sand in a growth chamber with the same regime described before and watered with a modified Long-Ashton (LA) solution containing 3.2 μM Na_2_HPO_4_·12H_2_O.

Mycorrhizal roots were collected at two-time points: 10 days post inoculation (dpi), and 40 dpi corresponding to the early and later stages of the mycorrhization process. For the molecular analyses, roots were immediately frozen in liquid nitrogen and stored at −80 °C. At the last time point (40 dpi), mycorrhizal roots were stained with cotton blue (0.1% in lactic acid), and the mycorrhizal colonization level was determined according to Trouvelot et al. (Trouvelot et al., 1986).

### Transcript analysis of mycorrhizal plants

Total RNA was extracted from rice roots using the Qiagen Plant RNeasy Kit according to the manufacturer’s instructions (Qiagen, Hilden; Germany). Following the producer’s instructions, samples were treated with TURBO™ DNase (ThermoFischer). The RNA samples were routinely checked for DNA contamination through PCR analysis. Single-strand cDNA was synthesized from 1 μg of total RNA using Super-Script II (Invitrogen), according to the instructions in the user manual. Quantitative RT-PCR (qRT-PCR) was performed using a Rotor-Gene Q 5plex HRM Platform (Qiagen). Each reaction was carried out in a total volume of 15 μL containing 2 μL of diluted cDNA (about 10 ng), 7.5 μl of 2× SYBR Green Reaction Mix, and 2.75 μl of each primer (3 μM). The following PCR program was used: 95°C for 90 s, 40 cycles of 95°C for 15 s, and 60°C for 30 s. A melting curve (80 steps with a heating rate of 0.5°C per 10 s and a continuous fluorescence measurement) was recorded at the end of each run to exclude the generation of non-specific PCR products.

All reactions were performed on at least three biological and two technical replicates. Baseline range and take-off values were automatically calculated using Rotor-Gene Q 5plex software.

The transcript level of *OsPt11* (an AM marker gene) was normalized using the *OsRubQ1* housekeeping gene (Güimil et al., 2005). Only take-off values leading to a mean with a standard deviation below 0.5 were considered. Statistical elaborations were performed using PAST statistical (version 4) (Hammer et al., 2001).

### Ethynyl deoxyuridine (EdU) staining for cell proliferation analysis

For the EdU staining, the seedlings were transferred to 50 ml falcon tubes containing 2 μM 5-ethynyl-2’-deoxyuridine (EdU) in dH_2_O, so that the roots were completely submerged in the solution, and kept there for 2 hr. The EdU staining was performed as decribed previously, using the Click-iT EdU Alexa Fluor 647 Imaging Kit (Invitrogen, ThermoFisher scientific, USA) (Kirschner et al., 2017). The root were cleared in CLEARSEE clearing solution (Kurihara et al., 2015) at 4 °C in darkness for two weeks, and cell walls were counterstained with 0.1 % Calcofluor White M2R in CLEARSEE overnight in darkness. After washing the roots in CLEARSEE, they were imaged using an inverted confocal microscope (LSM 710, Zeiss) and a 20x objective. Calcofluor White was excited with 405 nm and detected in a detection range of 410 -585 nm. Alexa 647 was excited with 633 nm and detected in a detection range of 638 – 755 nm.

## Statistical analysis

Data are represented as mean and their variations as standard deviation. The statistical significance was determined by one-way analysis of variance (one-way ANOVA) and Tukey’s multiple comparison test, using a probability level of p<0.05. All statistical elaborations were performed using GraphPad Prism 9.

## Data availability

All data needed to evaluate the conclusions in the paper are present in the paper and/or the Supplementary Materials. RNA-Seq data can be accessed at NCBI’s Gene Expression Omnibus (GEO) via accession number (GSE221837).

## Author Contributions

S.A.-B. and J.Y.W. proposed the concept. J.Y.W and G.-T. E. C. designed the experiments. J.B. generated transgenic *Osmax1* mutants. J.B. and G.-T. E. C. conducted genotyping. C.V., V.F., and L.L. investigated mycorrhization studies. J.Y.W. and G.-T. E. C. performed LC-MS analysis as well as RNAseq sample preparation and data analysis. G.-T. E. C., G.K.K., and I.B. prepared and performed cellular level analysis. M.J., J.Y.W., and G.-T. E. C. conducted Striga bioassays. G.-T. E. C., J.Y.W., J.B., M.J., L.B., and A.B. conducted phenotyping experiments. G.-T. E. C., J.Y.W., I.B., L.L., and S. A.-B. analyzed and discussed the data. G.-T. E. C., J.Y.W., J.B., and S. A.-B. wrote the original manuscript. All authors read, edited, and approved the manuscript.

## Acknowledgments

We sincerely thank the support from the members of KAUST Analytical Core Lab and the Bioactives lab. We thank Dr. Abdel Gabbar Babiker for providing *Striga hermonthica* seeds and Prof. Tadao Asami for providing TIS108. This work was supported by baseline funding given to S. A-B from King Abdullah University of Science and Technology (KAUST) and the Bill and Melinda Gates Foundation (grant number OPP1136424).

