## Supplemental Figures for "Disruption of the rice *4-DEOXYOROBANCHOL HYDROXYLASE* unravels specific functions of canonical strigolactones"

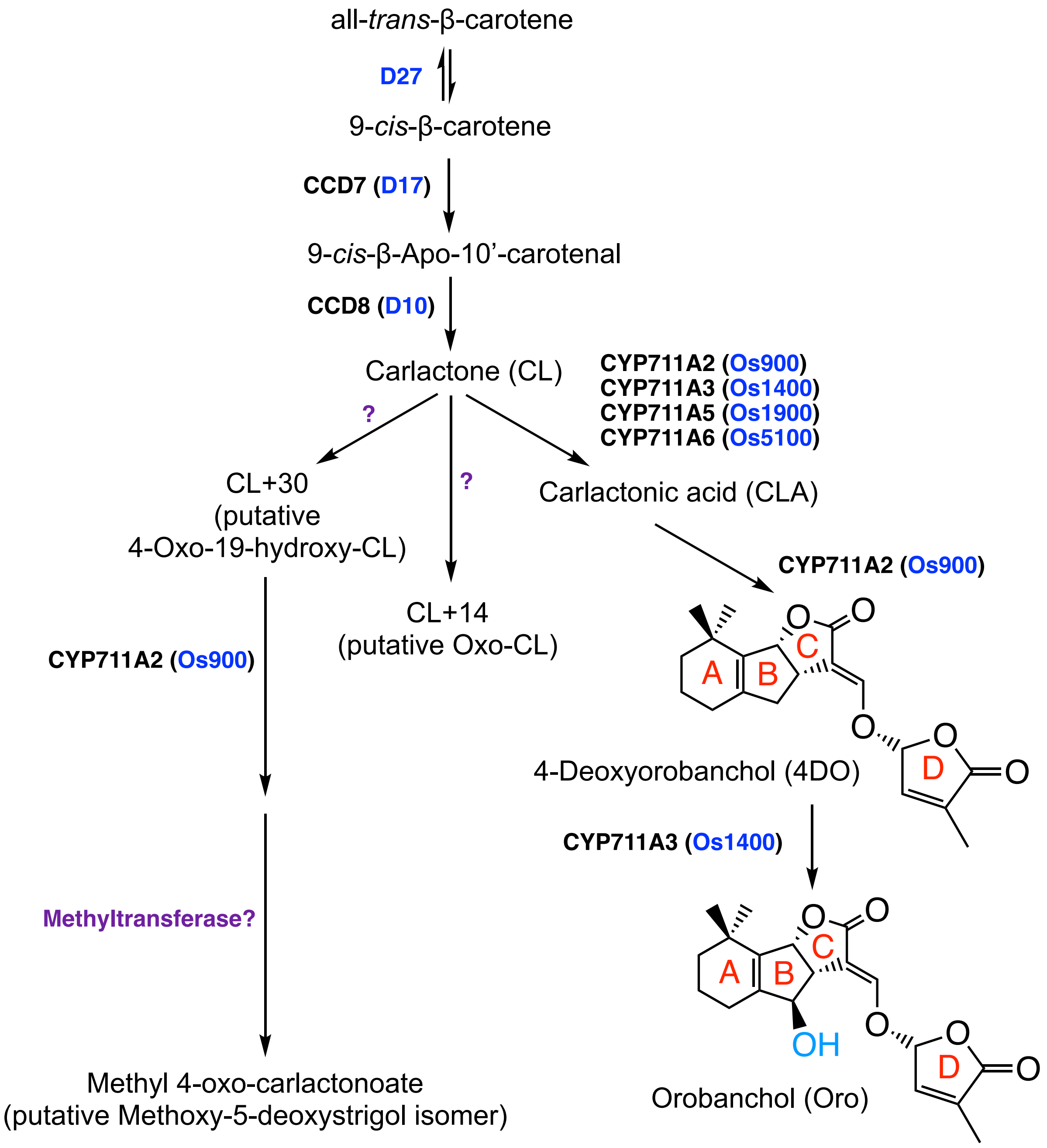


**Fig. S1. SL biosynthetic pathway in rice.**
Abbreviations: D27, Dwarf27; CCD, Carotenoid Cleavage Dioxygenase; MAX1, More Axillary Growth 1; CYP, Cytochrome P450

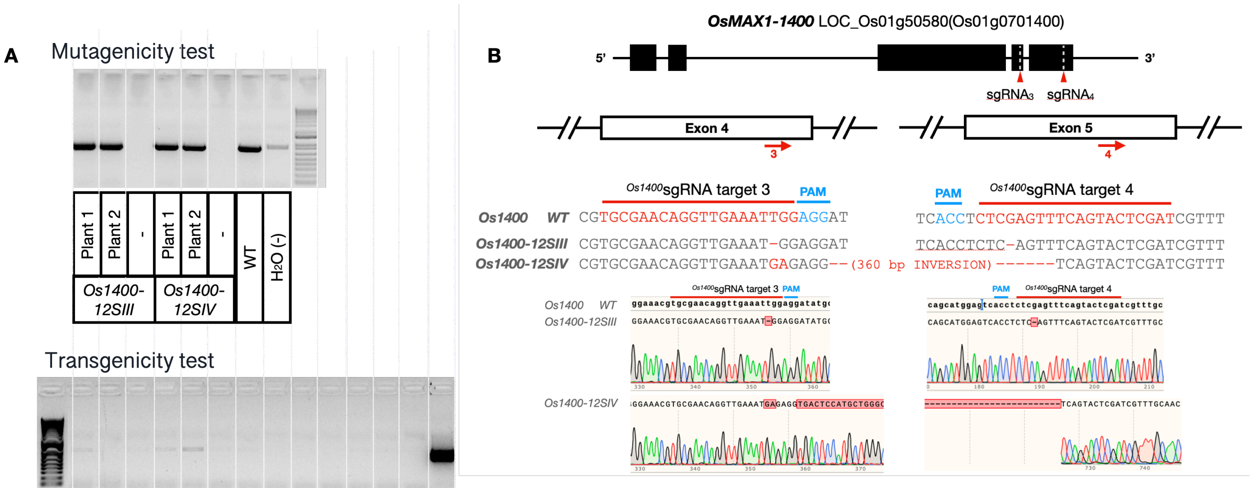


| Plant name | protein sequence at 1400-gRNA 3 | Protein size (aa) | Comments |
|---|---|---|---|
| WT | Cys Glu Gln Val Glu Ile Gly Gly Tyr Ala | 542 aa | |
| 12SIII-8 | Cys Glu Gln Val Glu Met Glu Asp Met Pro Phe .... * | 441 aa | Stop codon before the Heme |
| 12SIV-3 | Cys Glu Gln Val Glu Met Arg Gly Asp Ser Met .... * | 436 aa | Stop codon before the Heme with big insertion |

**Fig. S2. Genotyping of *Os1400*-KO lines.**

(A) Genomic DNA amplification of the region surrounding sgRNA target site in wild-type (WT) and *Os1400*-KO lines *12SIII* and *12SIV* (2 plants each) (up, mutagenicity test) and pRGEB32 region containing the two *Os1400* sgRNAs sequences (6 plants each) (down, transgenicity test). Water ($H_2O$) and the pRGEB32 vector containing the two *Os1400* sgRNAs sequences were used as a negative (-) and positive (+) control, respectively. (B) Sequencing details of two representative plants of the homozygous *Os1400*-KO lines showing the different mutations present in each line, aligned to the WT sequence for both *Os1400* sgRNAs target sites. (C) Prediction of protein sequences revealed a early stop codon before the heme-iron ligand signature that is necessary for P450 protein activity. Abbreviations: WT, wild-type.

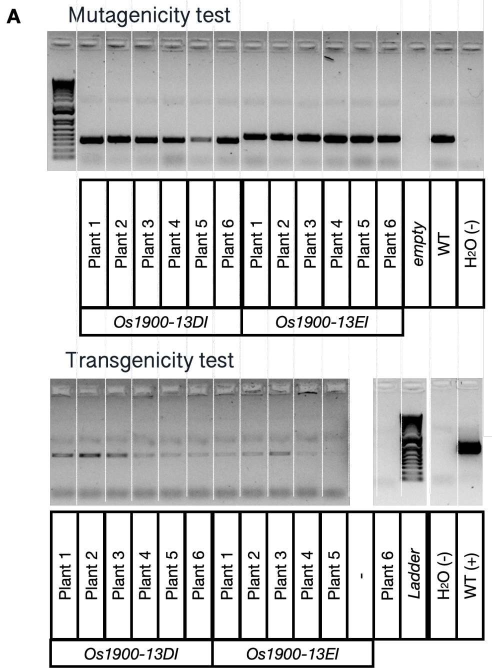


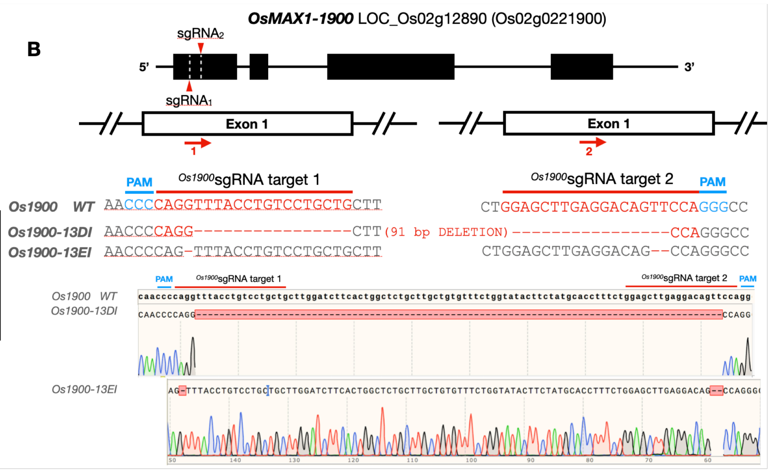


C

| Plant name | protein sequence at 1900-gRNA 1 | Protein size (aa) | Comments |
|---|---|---|---|
| WT | Pro Gly Leu Pro Val Leu Leu Leu Gly Ser Ser | 548 aa | |
| 13DI-8-2 | Pro Gly Gln Gly Leu .... * | 32 aa | Stop codon after transmembrane domain |
| 13EI-8-1 | Pro Val Tyr Leu Ser Cys Cys .... * | 50 aa | Stop codon after transmembrane domain |

**Fig. S3. Genotyping of *Os1900*-KO lines.**
(A) Genomic DNA amplification of the region surrounding sgRNA target site in wild-type (WT) and *Os1900*-KO lines *13DI* and *13EI* (6 plants each) (up, mutagenicity test) and pRGEB32 region containing the two *Os1900* sgRNAs sequences (down, transgenicity test). Water ($H_2O$) and the pRGEB32 vector containing the two *Os1900* sgRNAs sequences were used as a negative (-) and positive (+) control, respectively. (B) Sequencing details of two representative plants of the homozygous *Os1900*-KO lines showing the different mutations present in each line, aligned to the WT sequence for both *Os1900* sgRNAs target sites. (C) Prediction of protein sequences revealed a early stop codon before the heme-iron ligand signature that is necessary for P450 protein activity. Abbreviation: WT, wild-type.

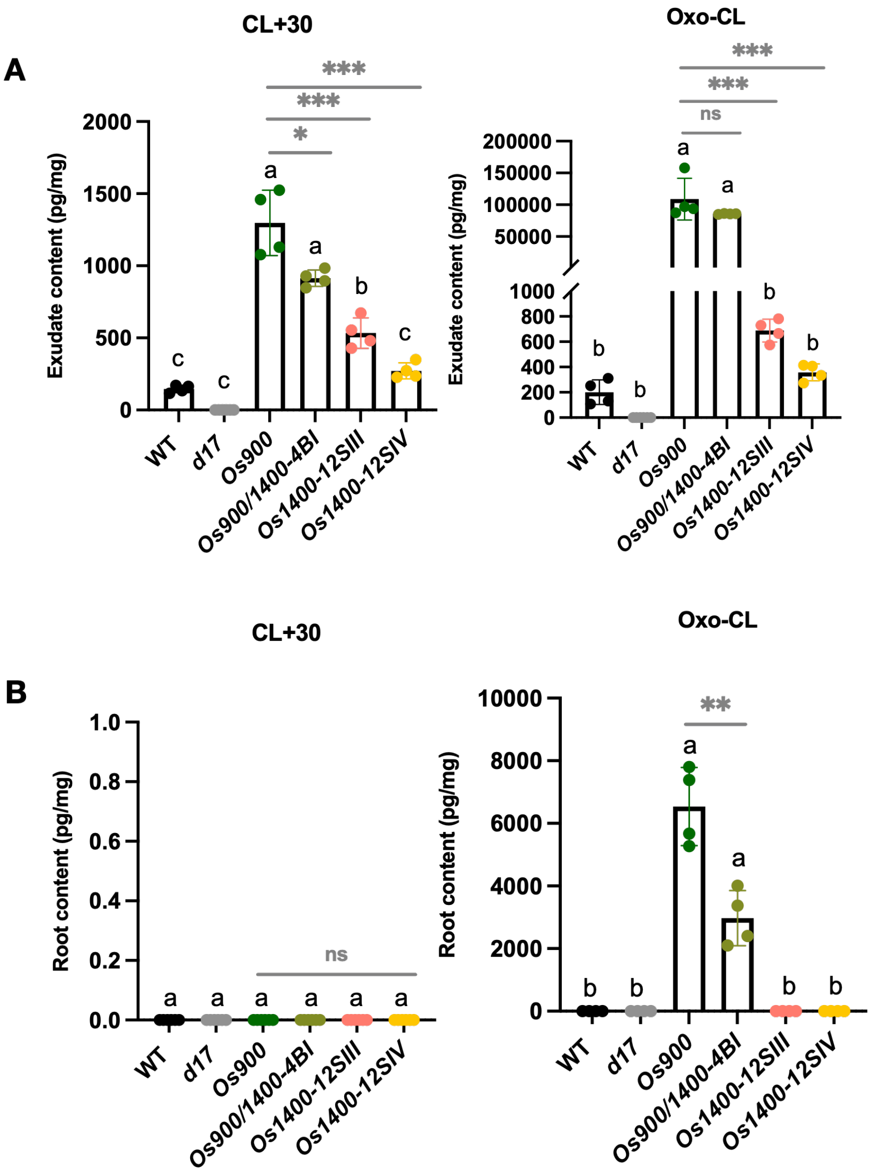


**Fig. S4. Quantification of putative non-canonical SLs in *Os900*-KO, *Os900/1400*-KO, and *Os1400*-KO lines**

LCMS quantification of SLs in the root exudates and root tissues of WT*, Os900-KO line, Os900/1400-KO line, Os1400-KO lines, and d17* mutant plants grown under low Pi conditions. Quantification of non-canonical SLs, CL+30 and oxo-CL, in (A) root exudates and (B) root tissues of WT*, Os900-KO line, Os900/1400-KO line, Os1400-KO lines, and d17* mutant plants grown under constant low Pi conditions. The data are presented as means ± SD for the number of biological replicates n=4 for (A) and (B). Significant values determined by one-way ANOVA are shown with different letter ($P < 0.05$) when compared to WT, and asterisks indicate statistically significant differences as compared to control by two tailed paired Student's t-test (*$p < 0.05$, **$p < 0.01$; ***$p < 0.001$; ****$p < 0.0001$). Abbreviations: CL, carlactone; WT, wild-type; ns, non-significant.

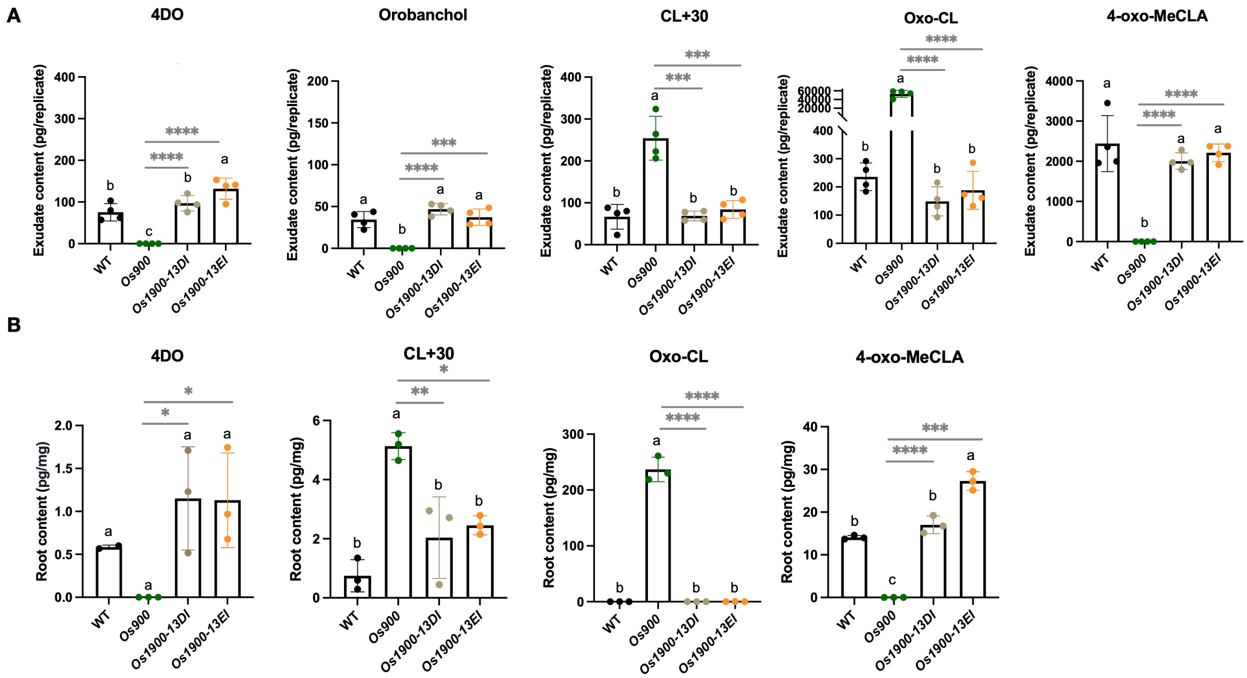


**Fig. S5. Quantification of different rice SLs for *Os900*-KO, and *Os1900*-KO lines**
(A) Analysis of SLs in the root exudates of WT, *Os900*-KO, and *Os1900*-KO mutant mutants grown in hydroponic culture under lowPi conditions. (B) Analysis of SLs in the root tissues of WT, *Os900*-KO, and *Os190*0-KO lines grown in hydroponic culture under lowPi conditions. The data are presented as means ± SD for the number of biological replicates n=4 for (A), and n=3 for (B). Significant values determined by one-way ANOVA are shown with different letter ($P < 0.05$) when compared to WT, and asterisks indicate statistically significant differences as compared to control by two tailed paired Student's t-test (*$p < 0.05$, **$p < 0.01$; ***$p < 0.001$; ****$p < 0.0001$). Abbreviations: CL, carlactone; 4- oxo-MeCLA, methyl 4-oxo-carlactonoate; WT, wild-type.

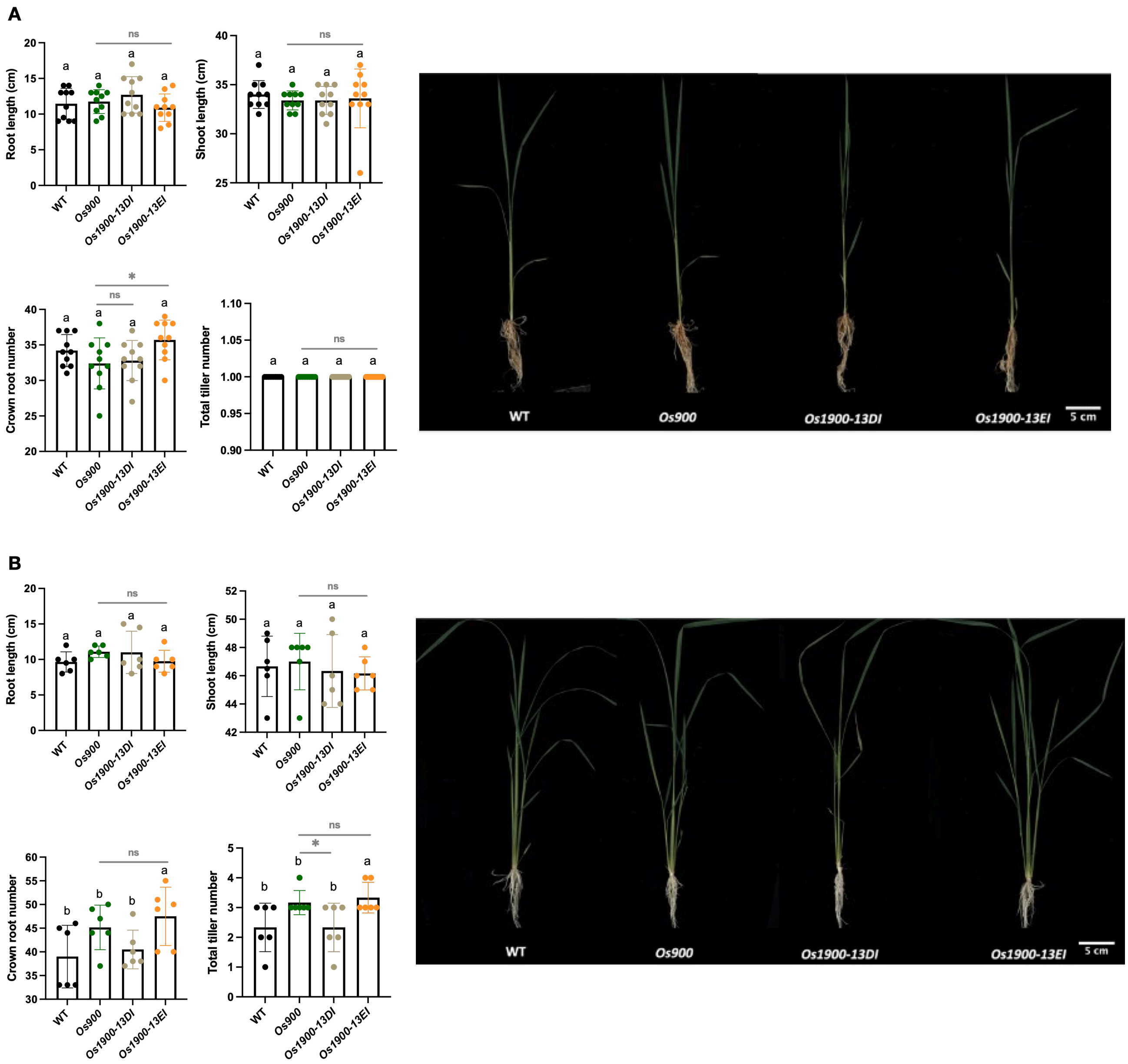


**Fig. S6. Phenotypic characterization of *Os900*-KO, and *Os1900*-KO lines**
Shoot and root phenotypes of WT, *Os900*-KO, and *Os1900*-KO mutant plants grown in hydroponic culture under (A) lowPi conditions and (B) normal (+Pi) conditions. The data are presented as means ± SD for the number of biological replicates n=10 for (A), and n=6 for (B). Significant values determined by one-way ANOVA are shown with different letter ($P < 0.05$) when compared to WT, and asterisks indicate statistically significant differences as compared to control by two tailed paired Student's t-test (*$p < 0.05$, **$p < 0.01$; ***$p < 0.001$; ****$p < 0.0001$). Scale bar, 5 cm. Abbreviations: WT, wild-type; ns, non-significant.

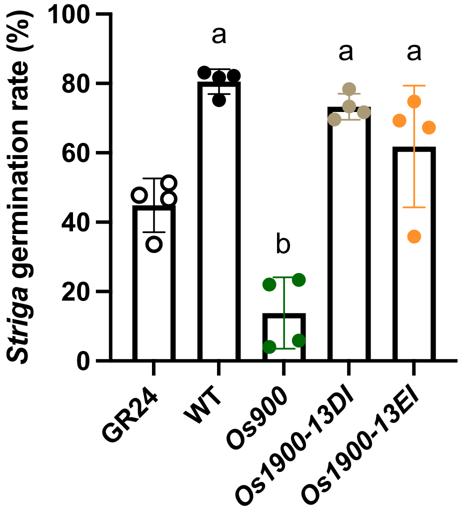


**Fig. S7. Striga germination test in the root exudates of *Os900*-KO, and *Os1900*-KO lines**. The data are presented as means ± SD for the number of biological replicates n=4. Significant values determined by one-way ANOVA are shown with different letter (P < 0.05) when compared to WT. Abbreviations: WT, wild-type.

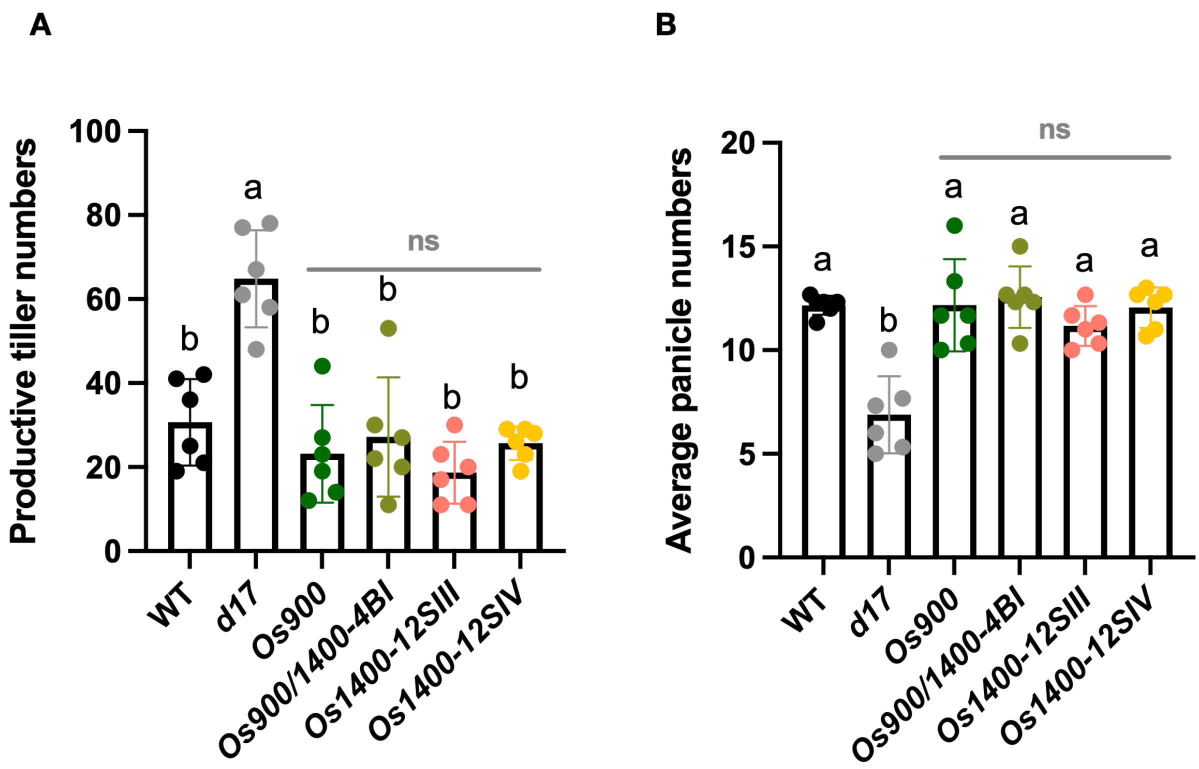


**Fig S8. Phenotypic characterization of of WT, *Os900*-KO, *Os900/1400*-KO, *Os1400*-KO, and *d17* mutant plants grown in soil.**
(A) Productive tiller numbers, and (B) average panicle numbers. The data are all presented as means ± SD for the number of biological replicates n=6. Significant values determined by one-way ANOVA are shown with different letter ($P < 0.05$) when compared to WT, and asterisks indicate statistically significant differences as compared to control by two tailed paired Student's t-test (*$p < 0.05$, **$p < 0.01$; ***$p < 0.001$; ****$p < 0.0001$). Abbreviations: WT, wild-type; ns, non-significant.

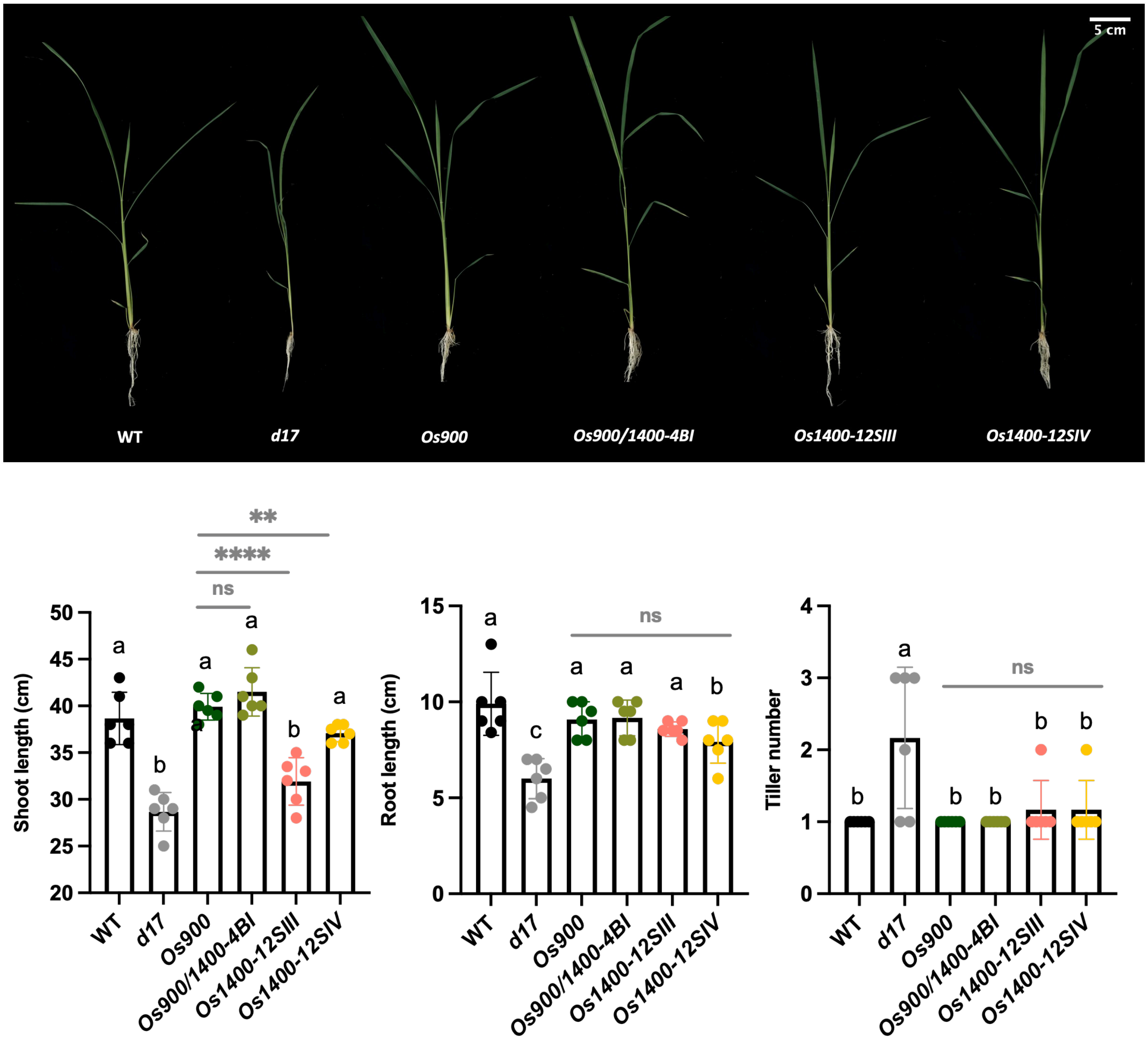


**Fig. S9. Shoot and root phenotypes of WT, *Os900*-KO, *Os900/1400*-KO, *Os1400*-KO, and *d17* mutants grown hydroponically under normal (+Pi) conditions.**
The data are all presented as means ± SD for the number of biological replicates n=6. Significant values determined by one-way ANOVA are shown with different letter ($P < 0.05$) when compared to WT, and asterisks indicate statistically significant differences as compared to control by two tailed paired Student's t-test (*$p < 0.05$, **$p < 0.01$; ***$p < 0.001$; ****$p < 0.0001$). Scale bar, 5 cm. Abbreviations: WT, wild-type; ns, non-significant.

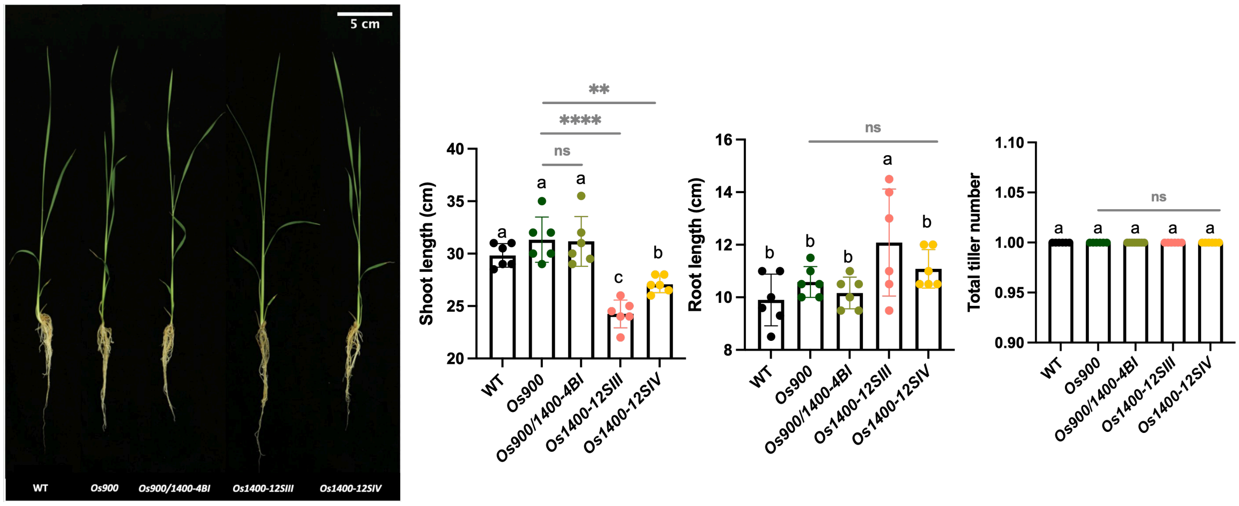


**Fig. S10. Shoot and root phenotypes of WT, *Os900*-KO, *Os900/1400*-KO, *Os1400*-KO, and *d17* mutants grown hydroponically under phosphate deficient (lowPi) conditions.** The data are all presented as means ± SD for the number of biological replicates n=6. Significant values determined by one-way ANOVA are shown with different letter ($P < 0.05$) when compared to WT, and asterisks indicate statistically significant differences as compared to control by two tailed paired Student's t-test (*$p < 0.05$, **$p < 0.01$; ***$p < 0.001$; ****$p < 0.0001$). Scale bar, 5 cm. Abbreviations: WT, wild-type; ns, non-significant.

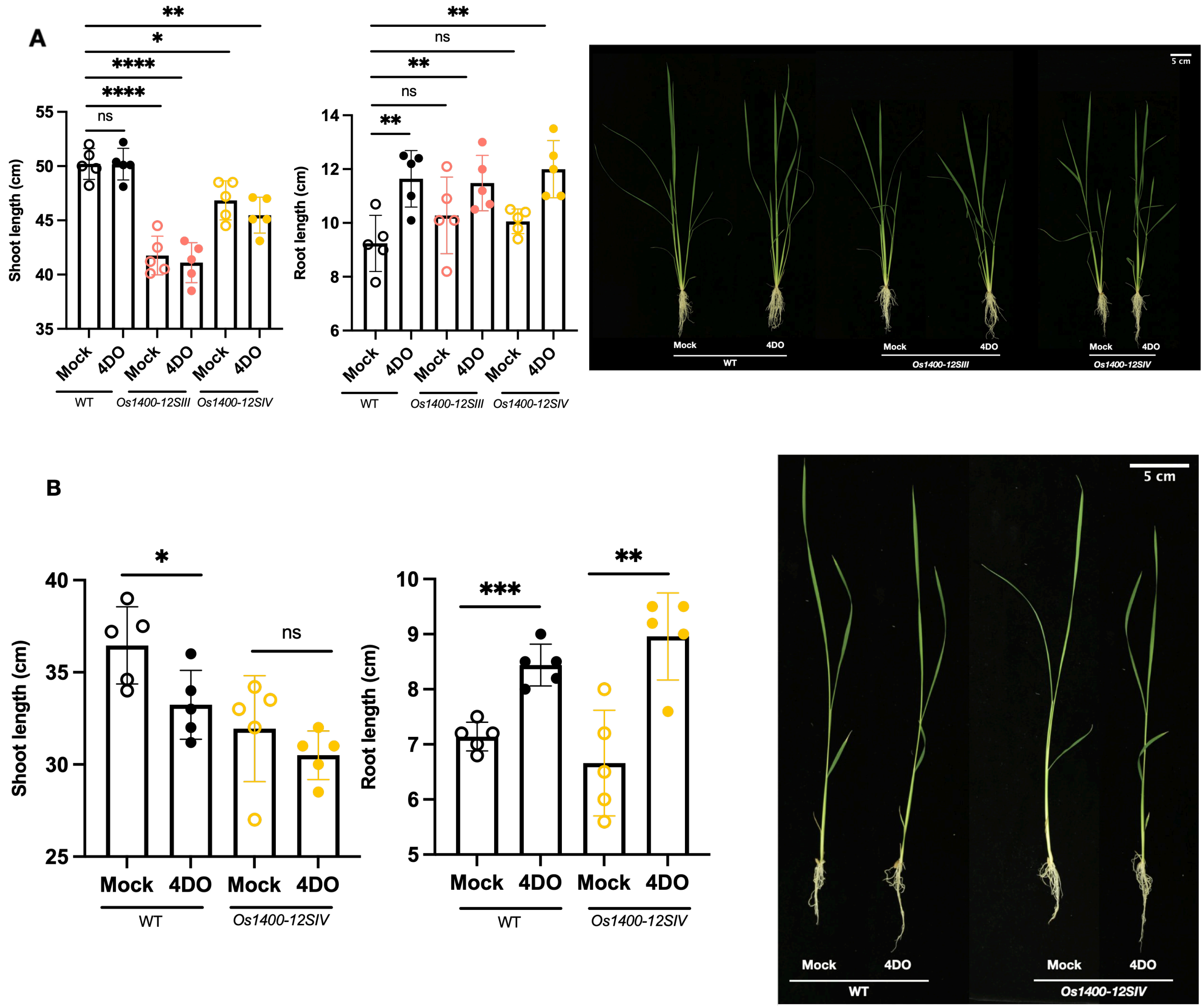


**Fig. S11. Effects of 4DO application on rice growth under normal (+Pi) hydroponic condition**
(A) Shoot and root phenotypes of WT, and *Os1400*-KO lines grown in hydroponic culture with or without (Mock) 300nM 4DO. (B) Shoot and root phenotypes of WT, and *Os1400*-KO lines grown in hydroponic culture with or without (Mock) 900nM 4DO. The data are all presented as means ± SD for the number of biological replicates n=5. Asterisks indicate statistically significant differences as compared to control by two tailed paired Student's t-test (*p < 0.05, **p < 0.01; ***p < 0.001; ****p < 0.0001). Scale bar, 5 cm. Abbreviation: 4DO, 4-deoxyorobanchol; WT, wild-type; ns, non-significant.

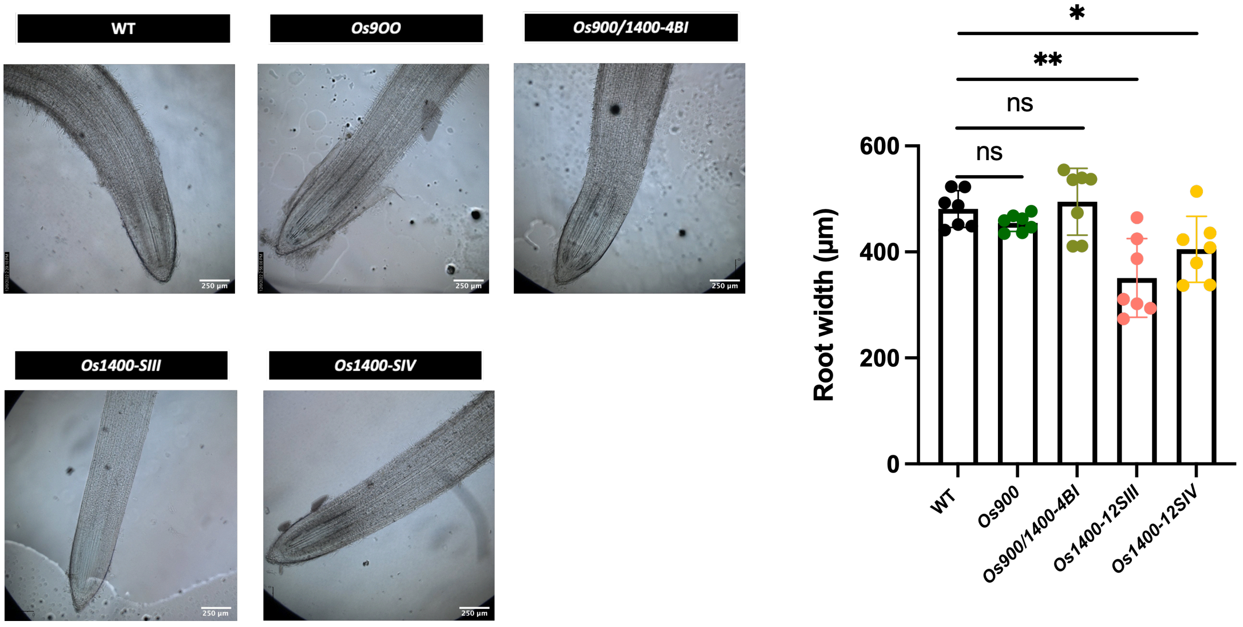


**Fig. S12. Root width (diameter) measurement of 2-week-old rice seedlings of Nipponbare WT, *Os900*-KO, *Os900/1400*-KO, and *Os1400*-KO lines.**
The data are all presented as means ± SD for the number of biological replicates n=7. Significant values determined by one-way ANOVA are shown with different letter (P < 0.05) when compared to WT, and asterisks indicate statistically significant differences as compared to control by two tailed paired Student's t-test (*p < 0.05, **p < 0.01; ***p < 0.001; ****p < 0.0001). Scale bar, 250 µm. Abbreviations: WT, wild-type; ns, non-significant.

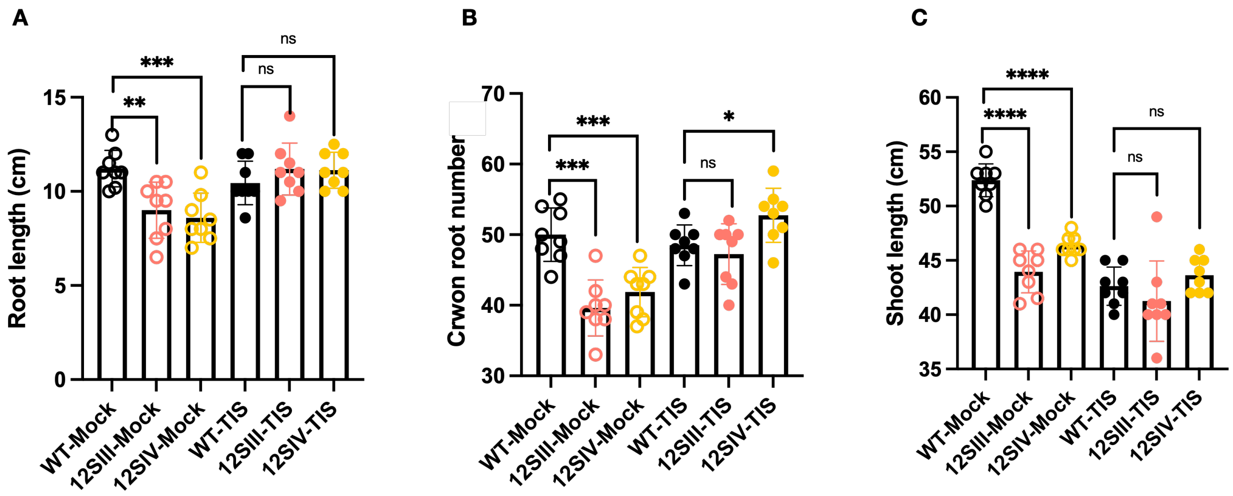


**Fig. S13. Effects of TIS108 on rice growth under normal (+Pi) hydroponic condition**

(A) Root length, (B) crown root numbers, and (C) shoot length. The data are all presented as means ± SD for the number of biological replicates n=8. Significant values determined by one-way ANOVA are shown with different letter ($P < 0.05$) when compared to WT, and asterisks indicate statistically significant differences as compared to control by two tailed paired Student's t-test (*$p < 0.05$, **$p < 0.01$; ***$p < 0.001$; ****$p < 0.0001$). Abbreviations: WT, wild-type; TIS, TIS108; ns, non-significant.

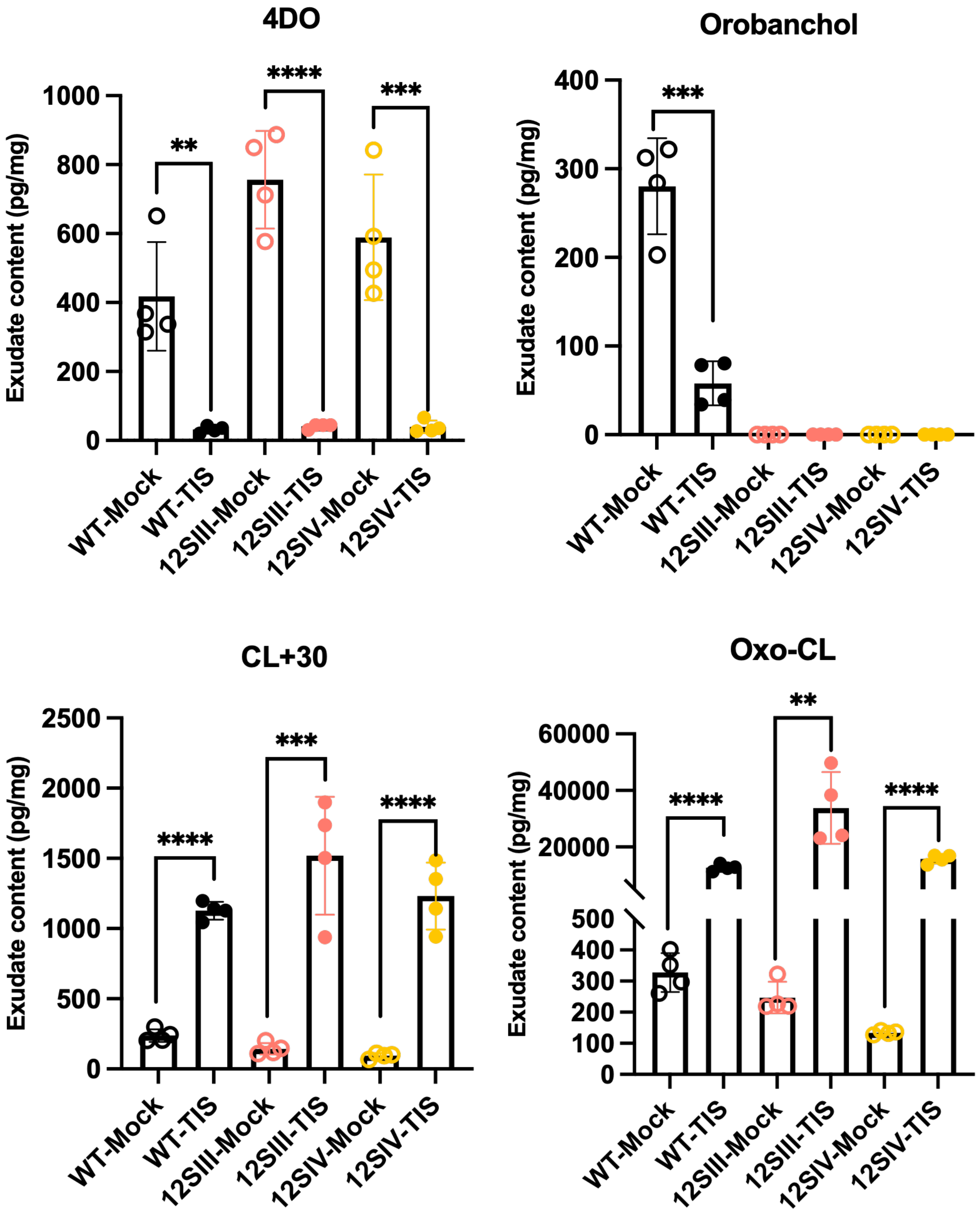


**Fig. S14. Effects of TIS08 on the quantification of different rice SLs in root exudates of WT and *Os1400* rice mutants.** The data are all presented as means ± SD for the number of biological replicates n=4. Asterisks indicate statistically significant differences as compared to control (Mock) within each line by two tailed paired Student's t-test (*$p < 0.05$, **$p < 0.01$; ***$p < 0.001$; ****$p < 0.0001$). Abbreviations: WT, wild-type; TIS, TIS108; ns, non-significant.

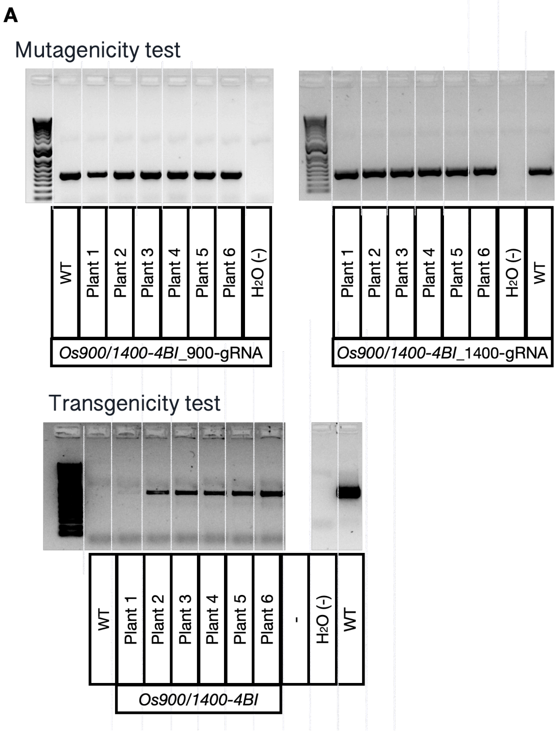


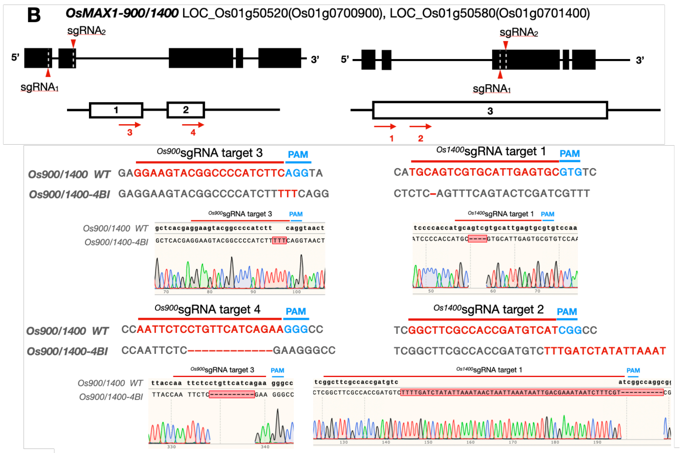


C

| Plant name | protein sequence at 900-gRNA 3 | Protein size (aa) | Comments |
|---|---|---|---|
| WT | Arg Lys Tyr Gly Pro Ile Phe Arg His Met | 540 aa | |
| 4BI-1-2 900(3,4) | Arg Lys Tyr Gly Pro Ile Phe Phe Arg .... | 135 aa | Early stop codon |

| Plant name | protein sequence at 1400-gRNA 1 | Protein size (aa) | Comments |
|---|---|---|---|
| WT | Met Gln Ser Cys Ile Glu Cys Val Ser Lys | 542 aa | |
| 4BI-1-2 1400(1,2) | Met Arg Ala Leu Ser Ala Cys Pro Arg .... | 170 aa | Early stop codon |

**Fig. S15. Genotyping of *Os900/1400*-KO lines.**
(A) Genomic DNA amplification of the region surrounding sgRNA target site in wild-type (WT) and *Os900/1400*-KO line *4BI* (6 plants) (up, mutagenicity test) and pRGEB32 region containing the two *Os900* sgRNA and *Os1400* sgRNA sequences (down, transgenicity test). Water ($H_2O$) and the pRGEB32 vector containing the two *Os900* sgRNA and *Os1400* sgRNA sequences were used as a negative (-) and positive (+) control, respectively. (B) Sequencing details of two representative plants of the homozygous *Os900/1400*-KO lines showing the different mutations present in each line, aligned to the WT sequence for both *Os900* sgRNA and *Os1400* sgRNA target sites. (C) Prediction of protein sequences revealed a early stop codon before the heme-iron ligand signature that is necessary for P450 protein activity. Abbreviations: WT, wild-type.

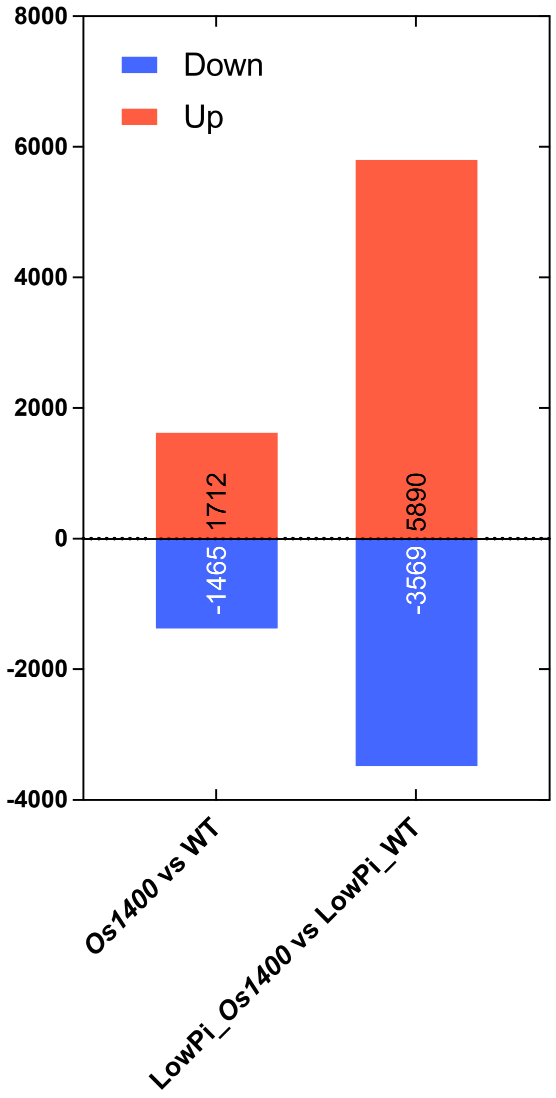


**Fig. S16. Differentially expressed genes (DEGs) under normal (+Pi) and phosphate deficient (lowPi) conditions.** Numbers of the significantly expressed genes (FDR < 0.05). Up- and down-regulated genes are shown in red and blue bars, respectively. Abbreviations: WT, wild-type.

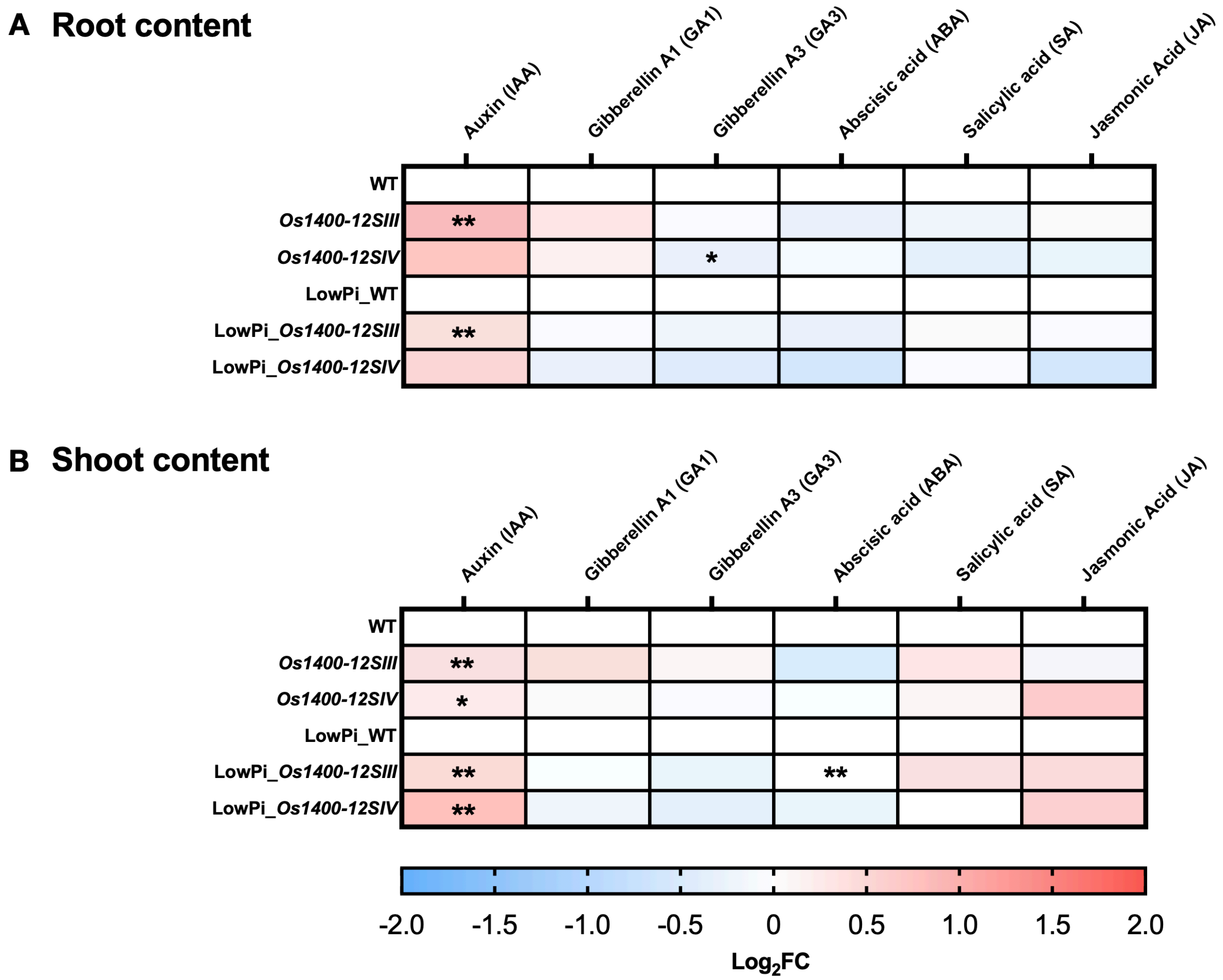


**Fig. S17. Hormone profile under normal (+Pi) and phosphate deficient (lowP) conditions.**

Heatmap showing relative accumulation of each hormone conten in (A) roots and (B) shoot bases (root-shoot junction) as compared to those in the WT. For each hormone, the value of the corresponding WT was set to 1. Asterisks indicate statistically significant differences as compared to control by t-test (*$p < 0.05$, **$p < 0.01$). Abbreviation: WT, wild-type.

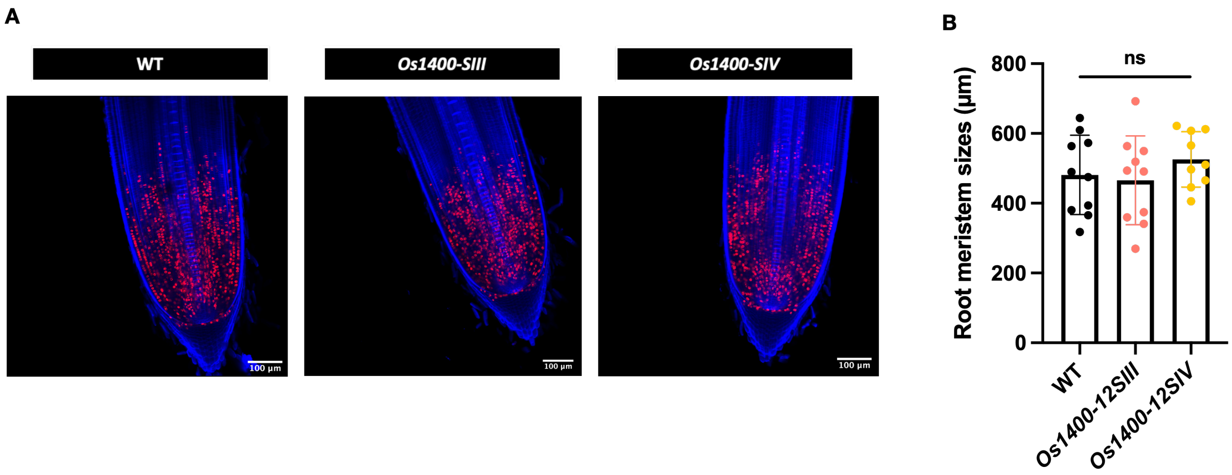


**Fig. S18 Characterization of root development of *Os1400*-KO lines at cellular level.**
(A and B) Ethynyl deoxyuridine (EdU) staining for cell proliferation analysis. Confocal images of rice root showing dividing cells as captured by EdU staining in Zeiss LSM 710 inverted confocal microscope. Root meristem length of 10-day-old rice seedlings of WT and *Os1400*-KO lines. Dividing EdU-stained nuclei are shown in red; cell walls counterstained with 0.1 % Calcofluor White M2R are shown in blue. Images were acquired using the tile scan function in the Zen software with automatized stitching. Regions of interest were divided into multiple tiles and imaged individually. The tiles were then combined via automatic stitching to create a large overview image. Images are representative of the total number ($n \geqslant 9$) of seedlings that were studied. The data are all presented as means ± SD for the number of biological replicates n=6. Significant values determined by one-way ANOVA are shown with different letter ($P < 0.05$) when compared to WT, and asterisks indicate statistically significant differences as compared to control by two tailed paired Student's t-test (*$p < 0.05$, **$p < 0.01$; ***$p < 0.001$; ****$p < 0.0001$). Scale bar, 100 µm. Abbreviations: WT, wild-type; ns, non-significant.

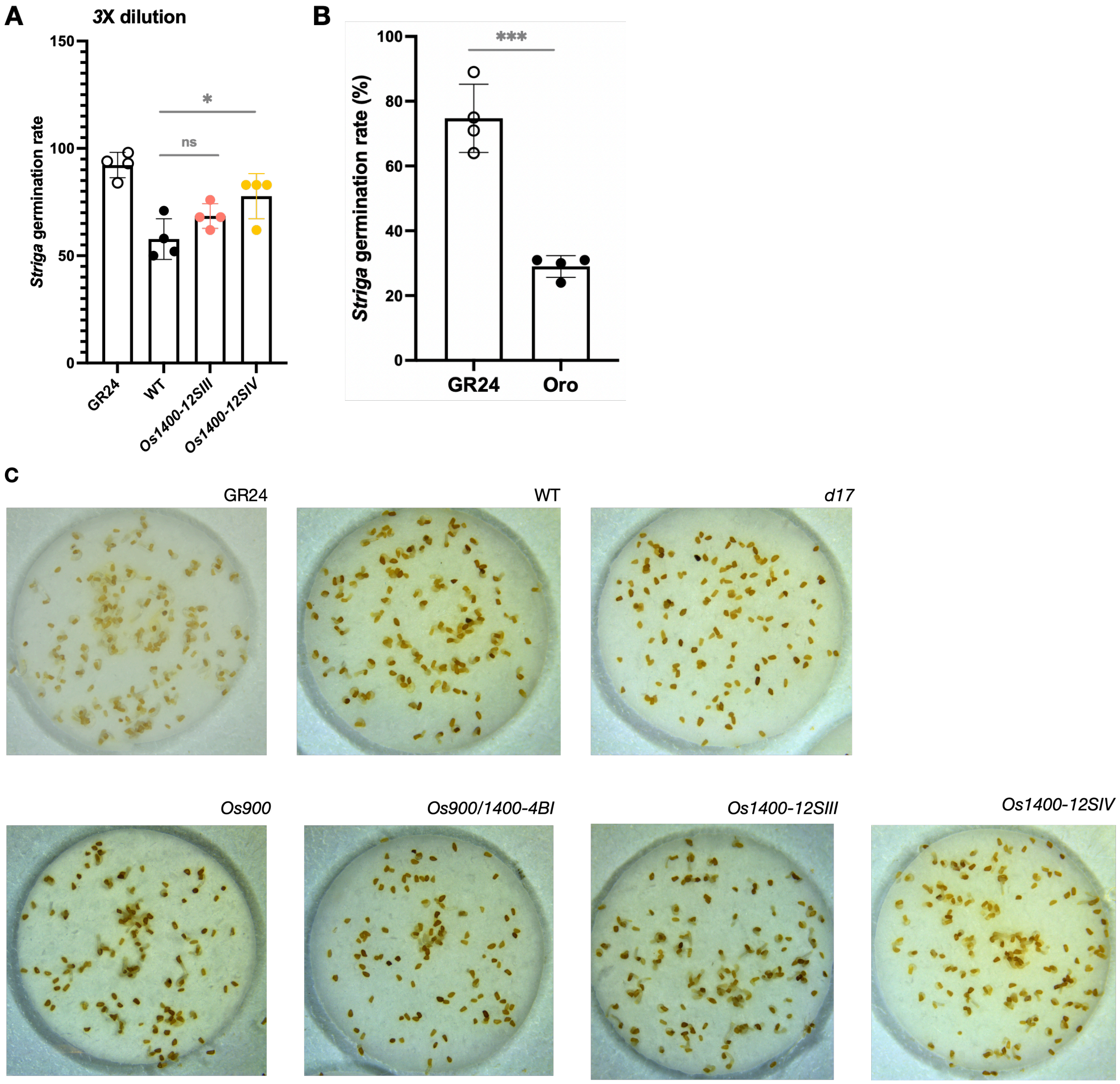


**Fig. S19. Striga seed germination assays.**
(A) Seed germination of root parasitic weeds (*Striga hermonthica*) by treatment with 3X dilution of the root exudates. (B) Seed germination of root parasitic weeds (*Striga hermonthica*) upon 1 μM GR24 and 10 μM Orobanchol application. (C) Seed germination of root parasitic weeds (*Striga hermonthica*) by treatment with 1X dilution of the root exudates. Significant values determined by two tailed paired Student's t-test (*p < 0.05, **p < 0.01; ***p < 0.001; ****p < 0.0001).Abbreviations: ns, non-significant.

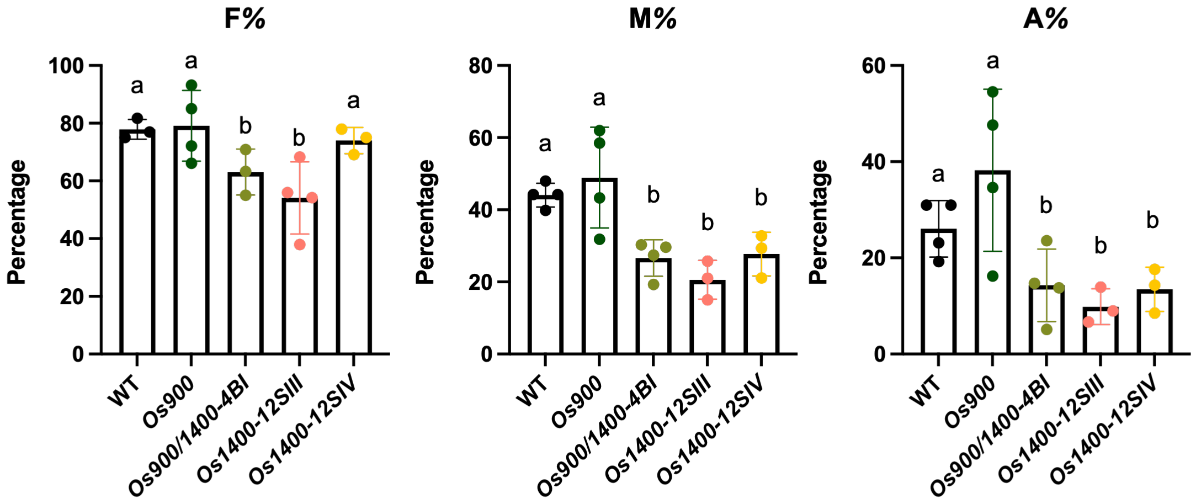


**Fig. S20. Evaluation AM symbiosis colonization in *Osmax1* mutant lines.** Mycorrhizal colonization of WT, *Os900/1400*-KO, and *Os1400*-KO lines by the AM fungus Rhizophagus irregularis at 40 dpi. Degree of colonization expressed as mycorrhizal frequency (F %), intensity (M %), and arbuscule abundance (A %) in the root system of WT and *Osmax1* lines. The data are all presented as means ± SD for the number of biological replicates n=4 for (A and B), and n≥3 for (C). Significant values determined by one-way ANOVA are shown with different letter ($P < 0.05$) when compared to WT. Abbreviations: WT, wild-type.
